# ATG9A and ARFIP2 cooperate to regulate PI4P levels for lysosomal repair

**DOI:** 10.1101/2024.07.23.604321

**Authors:** Stefano De Tito, Eugenia Almacellas, Daniel Dai Yu, Wenxin Zhang, Emily Millard, Javier H. Hervás, Enrica Pellegrino, Ioanna Panagi, Ditte Fodge, Theresa L.M Thurston, Maximiliano Gutierrez, Sharon A. Tooze

**Author notes:** These authors contributed equally (Stefano De Tito, Eugenia Almacellas).

## Abstract

Lysosome damage activates multiple pathways to prevent lysosome-dependent cell death, including a repair mechanism involving ER-lysosome membrane contact sites, phosphatidylinositol 4-kinase- 2a (PI4K2A), phosphatidylinositol-4 phosphate (PI4P) and oxysterol-binding protein-related proteins (ORPs), lipid transfer proteins. PI4K2A localizes to trans-Golgi network and endosomes yet how it is delivered to damaged lysosomes remains unknown. During acute sterile damage, and damage caused by intracellular bacteria, we show that ATG9A-containing vesicles perform a critical role in delivering PI4K2A to damaged lysosomes. ADP ribosylation factor interacting protein 2 (ARFIP2), a component of ATG9A vesicles, binds and sequesters PI4P on lysosomes, balancing ORP- dependent lipid transfer and promoting retrieval of ATG9A vesicles through recruitment of the adaptor protein complex-3 (AP-3). Our results reveal a role for mobilized ATG9A vesicles and ARFIP2 in lysosome homeostasis after damage and bacterial infection.

## INTRODUCTION

Lysosomal membrane permeabilization (LMP) is a process by which lysosomal membranes become leaky due to membrane damage. LMP is relevant in the context of neurodegenerative diseases, infection, and cancer^1–4^. To cope with LMP, several quality control mechanisms have been described to restore lysosome membrane integrity. Under physiological conditions, LMP can be constrained through lysosomal repair mediated by the endosomal sorting complex required for transport (ESCRT) machinery^5^. A phosphoinositide[initiated membrane tethering and lipid transport (PITT) pathway has also been described to support lysosomal repair^6^. Furthermore, larger lysosomal damage can recruit Annexin 1 and 2 ^7^ in a calcium-dependent and ESCRT-independent fashion to stem lysosomal leakage. ATG8s, in particular LC3A, can also be recruited and lipidated on damaged lysosomes together with ATG2 and independently of WIPI2/ATG13 to mediate lysosomal repair (11). Finally, the recruitment of stress granules is a mechanism for stabilizing the lysosomal membrane in response to damage utilizing ESCRT-dependent or independent pathways^8, 9^. Inadequate lysosomal repair leads to lysophagy, a selective form of macroautophagy triggered by the ubiquitination of lysosomal proteins^10^. In extreme cases, when damage is extensive, lysosome-dependent cell death is activated by the release of cathepsins into the cytosol^11^. How these pathways are regulated and coordinated to promote survival is still an outstanding open question.

In the PITT pathway, the phosphatidylinositol 4-kinase type 2a (PI4K2A) is recruited to lysosomes where it catalyzes the formation of phosphatidylinositol-4 phosphate (PI4P). PI4P, in turn, recruits, PI4P-binding proteins (ORP9, ORP10, ORP11 and OSBP) to establish ER-lysosome membrane contact sites which support lipid transfer for lysosomal repair^6, 12^. ATG2, a lipid transfer protein required for autophagy, is recruited to lysosomes upon damage and contributes to lysosomal repair^6, 13^. However, how PI4K2A is delivered and regulated on damaged lysosomes is still largely unknown.

ATG9A is a lipid scramblase whose function has been mainly studied in autophagy, specifically at the stage of phagophore formation and expansion, although under normal conditions it cycles between the trans-Golgi network (TGN) and endosomal compartments^14^. Under starvation conditions, ATG9A translocates in vesicles from the TGN to ER-proximal sites where ATG13, and remaining components of the ULK1 complex, are concomitantly recruited^15^. These ATG9A vesicles contain PI4KIIIβ, PI4K2A, and ARFIP2^16^. ARFIP2 is a Bin/Amphiphysin/Rvs (BAR)-domain-containing protein able to sense and induce membrane curvature^17, 18^. Through its BAR domain, ARFIP2 interacts with the small GTPases ARFs, ARL1, and RAC1^19–22^. Moreover, ARFIP2 possesses an amphipathic helix (AH), binding specifically to PI4P in *in vitro* membrane models, which, together with the BAR domain, is required for the correct localization of ARFIP2 at the TGN^23^. Further, ARFIP2 has been shown to coordinate the secretion of matrix metalloproteases 2 and 7, in complex with ARF1, ARL1 and PKD2^24^, and to positively regulate the autophagy pathway^16^.

ATG9A trafficking between different membrane compartments is coordinated by adaptor protein (AP) complexes^25^. ATG9A appears to be the main cargo of post-Golgi AP-4-coated vesicles, which are responsible for ATG9A trafficking to a vesicular compartment closely associated with autophagosomes^26^. AP-2 motifs, AP-2 binding to ATG9A and trafficking through the endosomal compartment has been shown to contribute to phagophore formation^27^. Recent studies have proposed a role for ATG9A in other membrane compartments, where, upon distinct stimuli, it contributes to alternative functions beyond autophagy, such as plasma membrane repair ^28^ lipid droplet homeostasis^29^ or Golgi function^30^. Interestingly, together with its localization at the TGN and early endosomes, ATG9A has been described more recently, to transiently interact with lysosomes^31, 32^. Intriguingly, and relevant to the PI4P and the PITT pathway, ATG9A-vesicles harbor PI4K2A^16^, and ATG9A interacts with ATG2 and modulates its lipid transfer activity *in vitro*^33, 34^. These facts prompted us to explore the potential involvement of ATG9A and the ATG9A-vesicle protein ARFIP2 in lysosomal repair mechanisms.

## RESULTS

### ARFIP2 loss increases lysosomal repair through ATG9A lysosomal retention

Upon acute lysosomal damage by the lysosomotropic agent L-Leucyl-L-leucine methyl ester (LLOMe), ATG9A traffics from the TGN, where it primarily resides in basal conditions, to lysosomes (Fig. 1a-b, Extended Data Fig. 1a-b). As autophagy induction triggers ATG9A dispersal from the TGN, we tested whether acute lysosomal damage might trigger autophagy and therefore mobilize ATG9A. Lysosome damage has been shown to inactivate the mammalian Target of Rapamycin complex 1 (mTORC1)^8^, which in turn inhibits macroautophagy, so we tested its activity by measuring the phosphorylation of S6K (T389), a downstream target. Importantly, mTORC1 activity was not affected after 15 minutes of LLOMe but inhibited at later time points (Fig. 1c, Extended Data Fig. 1c). As ATG13 translocates within seconds to ATG9A-positive structures ^15^ it is unlikely that mTORC1-inactivation dependent ULK1 activation is required. Importantly, LLOMe treatment did not significantly affect ATG9A levels (Extended Data Fig. 1d-e). LC3B lipidation, a hallmark of autophagy, was detectable by 15 minutes after LLOMe treatment and its levels drastically increased after 45 minutes in parallel to the reduction in mTORC1 activity (Fig. 1c-d). However, upon lysosomal damage, LC3B lipidation might be induced by the conjugation of ATG8s to single membranes (CASM), a pathway that is independent of canonical autophagy^13^. Thus, we used WIPI2 as a *bona fide* early autophagy marker independent of CASM. After 15 minutes of LLOMe treatment, the number of WIPI2 spots did not increase, ruling out autophagy initiation as the trigger for ATG9A mobilization (Fig. 1e-f). LC3B lipidation on lysosomes can trigger the association of ATG2 for lysosomal repair ^13^. As ATG9A interacts with ATG2, we tested whether ATG8s were indeed essential for ATG9A recruitment upon damage. To do so, we took advantage of the HeLa Hexa KO cell line, where all ATG8s have been knocked out^35^. Interestingly, ATG9A recruitment to lysosomes upon acute damage occurs in the absence of ATG8s (Fig. 1g-h), suggesting that its recruitment is independent of CASM.

**Fig. 1.**
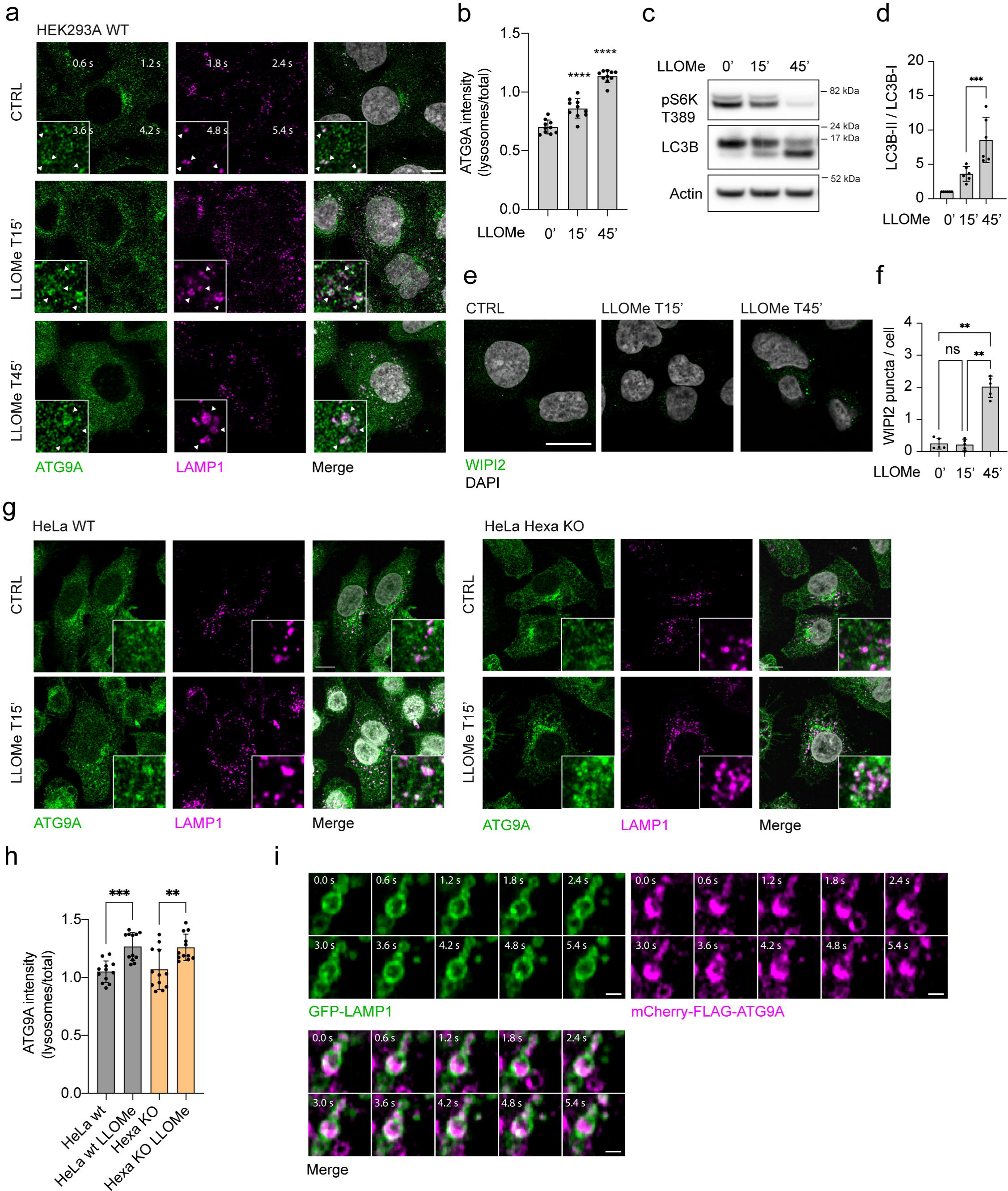
ATG9A is recruited to lysosome upon lysosomal damage. (a) HEK293A cells WT (CTRL) were treated with 1 mM LLOMe for the indicated times followed by immunofluorescence. Scale bar: 10 μm. (b) Quantification of ATG9A on the lysosomal compartment. *n* = 5 independent experiments, **** p < 0.0001. (c) HEK293A WT cells were treated with 1 mM LLOMe for the indicated times followed by Western Blot. (d) Quantification of LC3-II/LC3-I intensities ratio. *n = 6* independent experiments, *** p < 0.001. (e) HEK293A WT cells were treated with 1 mM LLOMe for the indicated times followed by immunofluorescence. Scale bar: 10 μm. (f) WIPI2 spots per cell were quantified in *n* = 3 independent experiments, ** p < 0.01. (g) HeLa WT and HeLa Hexa KO cells were treated with 1 mM LLOMe for 15 minutes followed by immunofluorescence. Scale bar: 10 μm (h) Quantification of ATG9A on the lysosomal compartment. *n* = 3 independent experiments, ** p < 0.01, *** p < 0.001. (i) HEK293A were transfected with GFP-LAMP1 and mCherry-FLAG-ATG9A and followed by live imaging. Scale bar: 1 μm.

Thus, we tested whether ATG9A vesicles can fuse with damaged lysosomes. To this aim, we performed live imaging experiments by over-expressing mCherry-ATG9A and GFP-LAMP1 in HEK293A cells that were treated with LLOMe. While the vast majority of ATG9A vesicles transiently interact with the lysosomes, a pool of ATG9A was detected on larger lysosomes forming a ring structure suggesting that fusion and fission events between ATG9A vesicles and the lysosomes may occur but at a low rate or very quickly, respectively (Fig. 1i, Movie S1). Due to the relevance of lysosomal integrity in neurons^36^, we tested if ATG9A lysosomal translocation upon damage was also occurring in a more physiological model. We used SH-SY5Y neuroblastoma cells and differentiated them into neuron-like cells prior to the treatment with LLOMe (Extended Data Fig. 1f-g) and confirmed ATG9A mobilization to lysosomes upon damage (Extended Data Fig. 1h).

A dispersal of ATG9A from its TGN localization was previously described upon loss of ARFIP2^16^. Thus, we tested whether ARFIP2 might regulate ATG9A localization on lysosomes. Interestingly, an enrichment of ATG9A in the lysosomal compartment was detected in HEK293A CrARFIP2KO cells compared to CTRL cells, which was further enhanced upon LLOMe treatment (Fig. 2a-b). Notably, acute LLOMe treatment did not affect ARFIP2 total protein levels (Extended Data Fig. 2a-b). We then investigated whether the lysosomal recruitment of ATG9A in CrARFIP2KO cells might impact lysosomal damage or repair. Galectin-3 (LGALS3) accumulation was used to detect damage to lysosomal membranes. Upon LLOMe treatment, CrARFIP2KO cells exhibited fewer LGALS3 spots than CTRL cells and this effect increased with the over-expression of GFP-ARFIP2 (CrARFIP2KO- rescue) (Fig. 2c-d, Extended data Fig. 2c-d). To test whether this phenotype depends on ATG9A on the lysosomes, we silenced ATG9A in either CTRL or CrARFIP2KO cells and analyzed LGALS3 spots upon LLOMe treatment. An increase of LGALS3 spots in the CrARFIP2KO without ATG9A suggested increased lysosomal damage (Fig. 2e-f, Extended Data Fig. 2e). As ARFIP2 interacts with PI4P via its AH, we tested whether the effects observed upon damage were dependent on the association of ARFIP2 to PI4P. We used the ARFIP2 W99A mutant of the AH, which loses the ability to associate with PI4P ^23^ and found upon damage, ARFIP2 W99A was unable to restore lysosomal repair, suggesting that ARFIP2 binding to PI4P is required for its function in lysosomal damage (Fig. 2c-d). To distinguish lysosomal damage from repair, we analyzed LysoTracker fluorescence recovery after LLOMe treatment. Lysotracker fluorescence requires lysosomal acidification, therefore decreased fluorescence reflects lysosomal damage which can be restored after repair^37^. Fluorescence recovery upon LLOMe-washout occurred faster and to a greater extent in CrARFIP2KO cells than CTRL or GFP-ARFIP2 rescued cells (Fig. 2g, Extended Data Fig. 2f). These data suggest that lysosomal repair is regulated by ARFIP2-dependent ATG9A accumulation to the lysosomes.

**Fig. 2.**
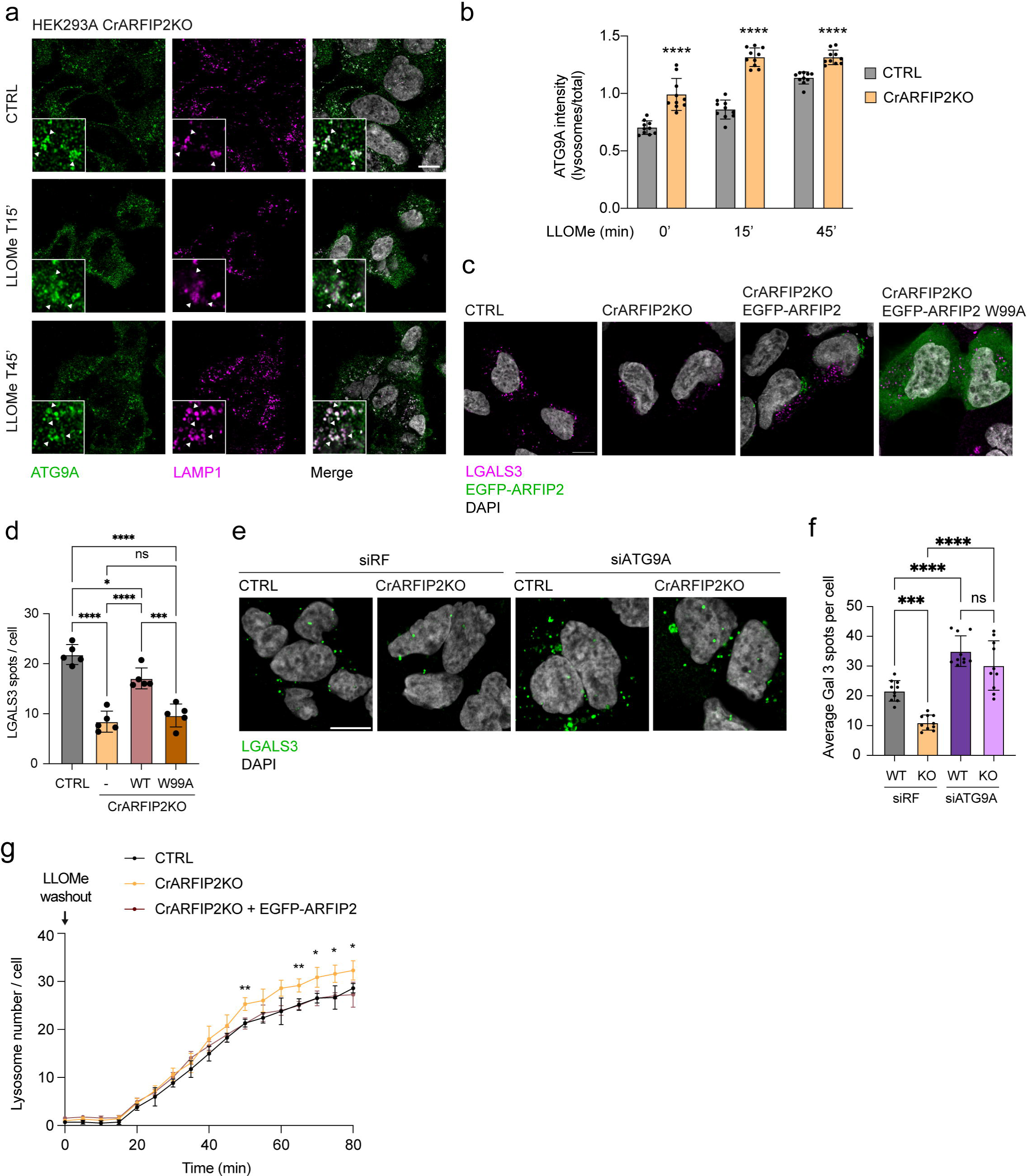
ARFIP2 controls ATG9A lysosomal localization for lysosomal damage. (a) HEK293A CrARFIP2KO cells were treated with 1 mM LLOMe for the indicated times followed by immunofluorescence. Scale bar: 10 μm. (b) Quantification of ATG9A on the lysosomal compartment. *n* = 5 independent experiments, **** p < 0.0001. (c) Control (CTRL) or CrARFIP2KO cell lines were transfected with EGFP-ARFIP2 or EGFP-ARFIP2^W99A^ mutant GFP- ARFIP2, treated with LLOMe 1 mM for 15 minutes followed by immunofluorescence. LGALS3 was used as a marker of lysosome damage. Scale bar: 10 μm. (d) Quantification of LGALS3 spots in (c). *n* = 5 independent experiments, * p<0.05 **** p < 0.0001. (e) Risc-free (siRF) or ATG9A siRNA was transfected into CTRL or CrARFIP2KO cells. After 72 hours, cells were treated with 1 mM LLOMe for 15 minutes and LGALS3 spots were detected by immunofluorescence. Scale bar: 10 μm. (f) Quantification of LGALS3 spots in CTRL and CrARFIP2KO after depletion of ATG9A. *n* = 5 independent experiments, *** p<0.001 **** p < 0.0001. (g) CTRL, CrARFIP2KO and rescue cells were loaded with 25 nM Lysotracker DND-99 for 45 minutes to stain the lysosomal compartment. Cells were subsequently treated with 1 mM LLOMe for 15 minutes, washed out and the fluorescence recovery was measured. *n* = 3 independent experiments, * p < 0.05, ** p < 0.01.

### AP-3 is an ARFIP2-interactor that regulates ATG9A trafficking through the endolysosomal compartment

We then investigated the mechanism underlying ARFIP2-mediated localization of ATG9A to lysosomes. Although ARFIP2 has been described as a Golgi-resident protein that binds PI4P^23^, live cell imaging experiments showed that GFP-ARFIP2 also localizes on lysosomes positive for RFP- LAMP1 (Fig. 3a, Movie S2). Immunoprecipitation experiments followed by mass spectrometry in HEK293A cells stably expressing GFP-ARFIP2 revealed the presence of several lysosomal proteins and proteins belonging to the PITT pathway (Fig. 3b).

**Fig. 3.**
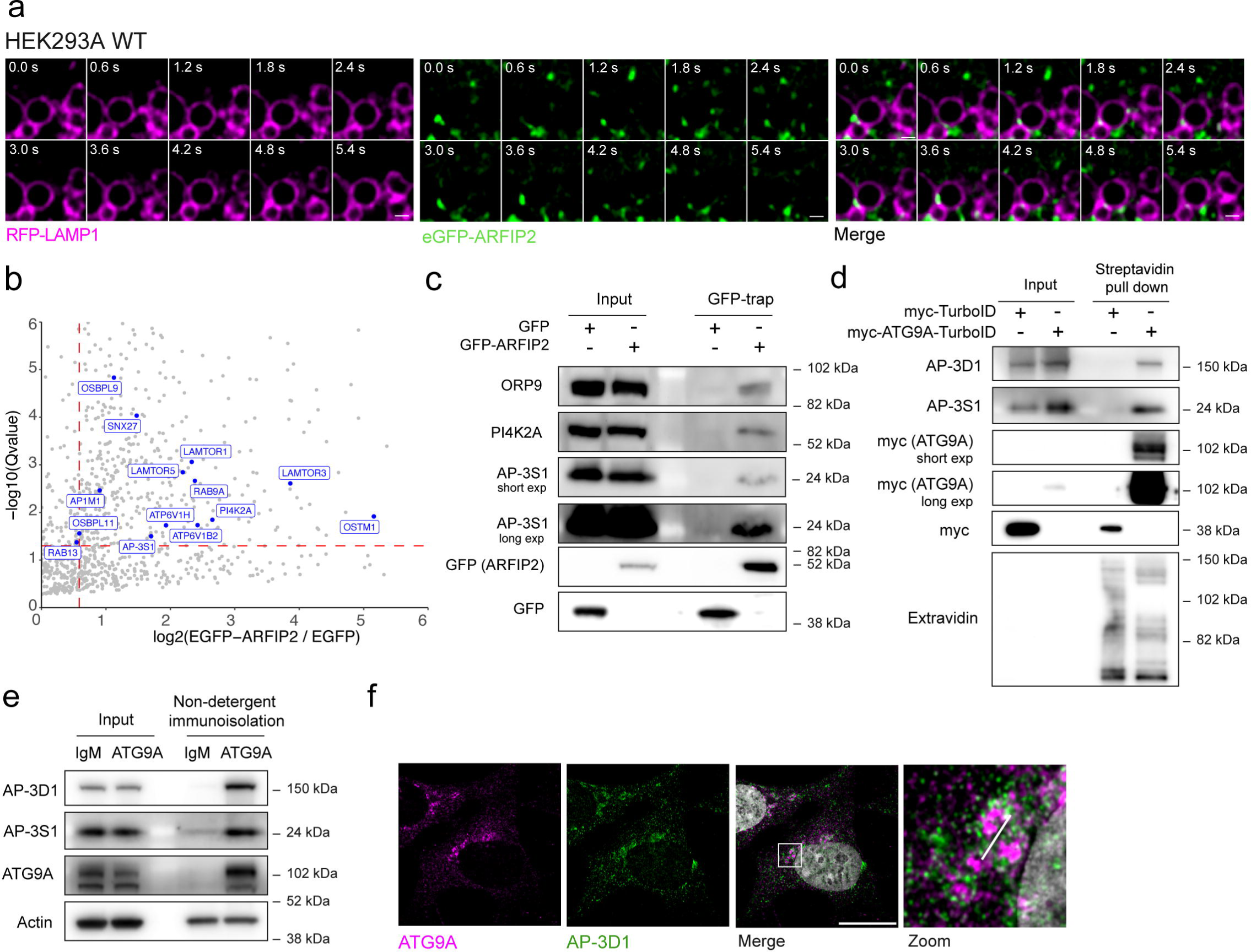
ARFIP2 interacts with the lysosomal compartment, lysosomal proteins and AP-3. (a) CrARFIP2KO cells were transiently transfected with eGFP-ARFIP2 (green) and RFP-LAMP (magenta). Frames from live imaging experiments were exported at different time points. Scale bar: 1 μm (b) GFP-trap was performed in CrARFIP2KO cells stably expressing GFP or GFP-ARFIP2. Label-free mass spectrometry analysis was performed. Lysosomal proteins (according to available data from Cell Map^69^) and proteins of the PITT pathway are highlighted in the volcano plot. (c) Validation of candidates was performed by GFP-trap followed by Western Blot analysis. (d) CrATG9AKO cells stably expressing myc-TurboID or myc-ATG9A-TurboID were treated with 50 μM biotin for 1 hour and subjected to streptavidin pulldown. Indicated proteins were detected by Western Blot. (e) Immunoisolation of endogenous ATG9A-positive membranes was performed in HEK293A WT cells. Indicated proteins were detected by Western Blot. (f) Immunofluorescence of ATG9A and AP-3D1 subunit was performed in HEK293A cells. Scale bar: 10 μm.

Among the ARFIP2-interactors, the adaptor complex (AP) AP-3 was detected (Fig. 3b-c). AP-3 is involved in the sorting of transmembrane proteins to/from the endolysosomal compartment^38–40^ Despite various AP complexes (AP-1, AP-2, AP-4) previously implicated in regulating ATG9A trafficking^26, 41–43^, a potential role for AP-3 was not known. ATG9A-proximal proteome generated from HEK293A cells stably expressing myc-ATG9A-TurboID mass spectrometry analysis, and western blot analysis revealed that two subunits of the adaptor protein complex AP-3 (AP-3B1 and AP-3S1) were enriched ATG9A interactors (Fig. 3d, Extended Data Fig. 3a-b). We confirmed that myc-ATG9A-TurboID colocalizes with and behaves like endogenous ATG9A, redistributing into peripheral cytosolic vesicles upon nutrient starvation (Extended Data Fig. 3c-d). Immunoisolation experiments of endogenous ATG9A-positive membranes further support this finding (Fig. 3e). Additionally, immunofluorescence analysis demonstrated similar intracellular localization of AP-3 and ATG9A to the Golgi complex (Pearson’s R coefficient: 0.684 ± 0.03209) (Fig. 3f). Since the depletion of a single subunit of an AP complex disrupts the formation and stability of the tetrameric complex^44^, we silenced the AP-3 sigma subunit (AP-3S1) in HEK293A cells and found this led to ATG9A dispersion (Extended Data Fig. 4a-c).

To validate the missorting of ATG9A upon loss of AP-3, we took advantage of fibroblasts from the *mocha* mouse model (MEF^mh/mh^), which are AP-3 deficient due to destabilization of AP-3D1 (Fig.4a)^45^. Genetic depletion of AP-3 has been described as the underlying genetic cause of Hermansky-Pudlak syndrome type 10 (HPS10), which is characterized by severe phenotypes such as inner ear degeneration, pigmentation dysfunctions and neurological deficits as pathological outcomes^45^. ATG9A localization differed between wild-type mouse embryonic fibroblasts (MEF^WT^) and MEF^mh/mh^, with the latter exhibiting accumulation of ATG9A in cytosolic vesicular structures (Fig. 4b-c). Importantly, overexpression of the AP-3D1 subunit in these cells rescued the expression of the full AP-3 complex as well as ATG9A localization (Fig. 4a-c). ATG9A dispersal in MEF^mh/mh^ cells was corroborated by differential centrifugation experiments (Extended Data Fig. 4d-f). Notably, comparing MEF^mh/mh^ with MEF^WT^ revealed enrichment of ATG9A in the lightest membrane fraction, in parallel with a decrease in heavier ones (Extended Data Fig. 4e-f). Interestingly, the levels of the lower band of ATG9A, corresponding to the ER-specific form^46^, were unchanged in the MEF^mh/mh^, suggesting that the AP-3 complex is not involved in ATG9A trafficking between the ER and the Golgi apparatus (Extended Data Fig. 4e). Live imaging experiments, using mCherry-3xFLAG- ATG9A and GFP-TGN46 to label the Golgi compartment in MEF^WT^ or MEF^mh/mh^ cells, provided dynamic insights into ATG9A constitutive trafficking. In contrast to MEF^WT^, ATG9A displayed a more dispersed localization barely colocalizing with the TGN in MEF^mh/mh^ cells, suggesting an impairment in the trafficking and/or recycling of ATG9A-positive vesicles (Fig. 4d, Movie S3, Movie S4).

**Fig. 4.**
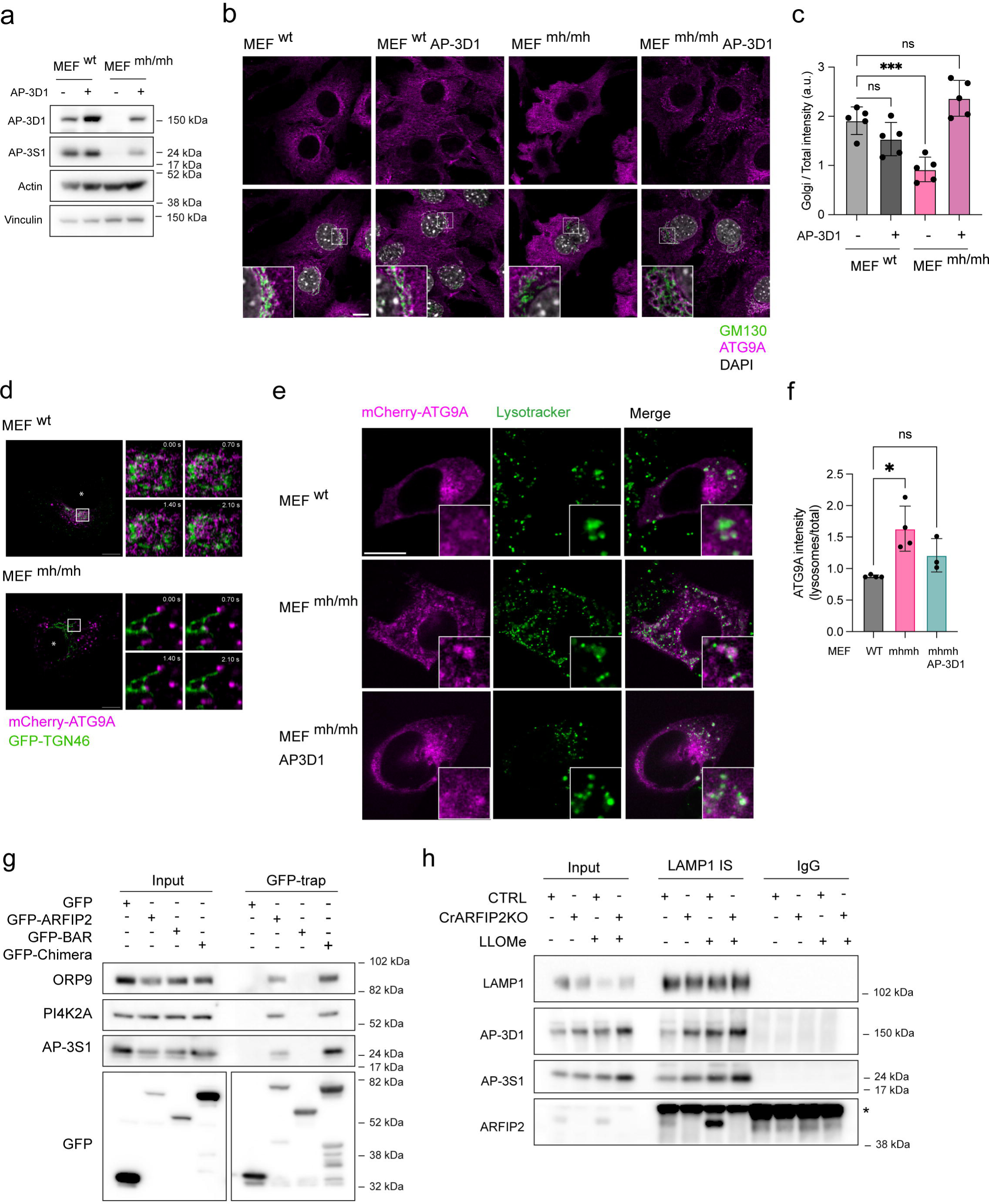
AP-3 controls ATG9A trafficking throughout the endolysosomal compartment. (a) MEF^wt^ and MEF^mh/mh^stably expressing AP-3D1 subunit were subjected to Western Blot. (b) Immunofluorescence staining of ATG9A and Golgi (GM130) in *wild type* mouse embryonic fibroblasts (MEF^wt^), MEF^wt^ stably expressing AP-3D1, MEF^mh/mh^ and MEF^mh/mh^ stably expressing AP-3D1. Scale bar: 10 μm. (c) Quantification of ATG9A in the Golgi in the indicated cell lines. *n* = 3 independent experiments, *** p < 0.001. (d) MEF^wt^ and MEF^mh/mh^ transiently expressing GFP- TGN46 and mCherry-FLAG-ATG9A were subjected to live imaging experiments. Frames were exported at the indicated times. Representative panels of *n* = 10 cells. (e) MEF^wt^, MEF^mh/mh^ and MEF^mh/mh^ stably expressing AP-3D1 were transiently transfected with mCherry-FLAG-ATG9A and loaded with Lysotracker Blue DND-26 (green) for one hour before live imaging. Scale bar: 10 μm (f) Quantification of ATG9A intensity overlapping with Lysotracker in overexpressing cells. *n = 4* fields were analyzed per condition, * p < 0.05 (g) CrARFIP2KO HEK293A cells stably expressing GFP, GFP-ARFIP2, GFP-AH-BAR (BAR) and GFP-ARFIP Chimera were subjected to GFP-trap. Indicated proteins were detected by Western Blot. (h) Lysosome immunoprecipitation using endogenous LAMP1 was performed in CTRL and CrARFIP2KO cells treated or not with LLOMe 1 mM for 15 minutes. Indicated proteins were detected by Western Blot.

In addition to the Golgi complex, ATG9A colocalizes with the endocytic-recycling compartment (ERC) and endosomes ^27, 47–52^ and partially with the lysosomal compartment^31, 32^, supporting the constitutive trafficking of ATG9A occurs throughout the endolysosomal system. Given the role of the AP-3 complex, we investigated whether ATG9A re-localized in the endolysosomal compartment in MEF^mh/mh^ cells. We loaded MEF^WT^ and MEF^mh/mh^ cells, expressing mCherry-3xFLAG-ATG9A, with fluorescent LysoTracker for 1 hour, to visualize the endolysosomal compartment (Fig. 4e-f). In live-cell imaging, the amount of ATG9A colocalizing with the endolysosomal compartment was remarkably increased in MEF^mh/mh^ compared to MEF^WT^ suggesting a defect in ATG9A sorting (Fig. 4e-f). Interestingly, cells re-expressing AP3 rescued this phenotype supporting a role for AP3 in ATG9A retrieval from the endolysosomal system (Fig. 4e-f). As ARFIP2 localizes to the lysosomal compartment and interacts with AP-3, these data support a model whereby ARFIP2 cooperates with AP-3 to retrieve ATG9A from lysosomes.

Next, we investigated which domain of ARFIP2 is involved in the interaction with AP-3S1. GFP-trap experiments show that ARFIP2 full-length, but not its AH-BAR (109-341) domain (GFP-BAR), interacts with AP-3S1 (Fig. 4g, Extended Data Fig. 4g). Differently from ARFIP2 AH-BAR-domain, the N-terminal ARFIP2-ΔBAR (1-108) domain does not retain its Golgi localization upon over- expression, hindering functional characterization (Extended Data Fig. 4g-h). As the AH-BAR domains of ARFIP1 and 2 are highly conserved^23^, and ARFIP1 is not involved in ATG9A vesicle trafficking^16^, we exploited this to test the role of the N-terminal domain of ARFIP2. We engineered an ARFIP Chimera (Extended Data Fig. 4g-h), by fusing the N-terminal portion of ARFIP2 (1-92) with the AH-BAR domain of ARFIP1 (94-341). Similar to ARFIP2, ARFIP Chimera localizes mainly in the Golgi and can interact with AP-3S1, while ARFIP2 AH-BAR domain did not, demonstrating that the N-terminal portion of ARFIP2 is responsible for its interaction with AP-3S1 (Fig. 4g). Surprisingly, this finding was consistent for two other ARFIP2 interactors, ORP9 and PI4K2A, involved in the PITT pathway (Fig. 4g). Finally, by immunoisolating LAMP-1 positive membranes, we also confirmed the presence of both ARFIP2 and AP-3 on the lysosomes and their enrichment upon lysosomal damage by treatment with LLOMe (Fig. 4h).

### ATG9A coordinates PI4K2A delivery to the lysosomes for lysosomal repair

AP-3 and PI4K2A interact, and both the dileucine motif and kinase activity of PI4K2A are required for AP-3 and PI4K2A localization to the lysosomal compartment^53–55^. Thus, we investigated whether loss of ARFIP2 affects the intracellular distribution of PI4K2A. Like ATG9A, PI4K2A was more dispersed and accumulated in the lysosomal compartment in CrARFIP2KO cells (Fig. 5a-b). Concomitantly, the colocalization between ATG9A and PI4K2A increase upon LLOMe treatment (Extended Data Fig. 5a-b). PI4K2A dispersal was supported by differential centrifugation experiments that showed enrichment of both PI4K2A and ATG9A in the lightest membrane fraction in CrARFIP2KO cells compared to CTRL cells (Extended Data Fig. 5c-e). Moreover, upon LLOMe treatment, PI4K2A accumulation was further enhanced on the lysosomes as shown by immunofluorescence and immunoisolation of LAMP1-positive membranes (Fig. 5a-b, Extended Data Fig. 5f). Considering that ATG9A membranes associate with PI4K2A^16^, we tested if PI4K2A mislocalization in CrARFIP2KO cells requires ATG9A (Fig. 5c-d). ATG9A siRNA knock-down (KD) induced an accumulation of PI4K2A in the peri-Golgi area not only in CrARFIP2KO cells but also in the CTRL cells, supporting the function of ATG9A-positive vesicles as carriers for the delivery of PI4K2A towards the endolysosomal compartment (Fig. 5c-d). PI4K2A is the main PI4P- producing enzyme at the lysosomes and has a prominent role in lysosomal repair^6, 12^. Indeed, the lysosomal accumulation of PI4K2A in CrARFIP2KO cells reflected an increased level of PI4P, as detected by the specific probe GFP-P4C-SidC ^56^ (Fig. 5e-f, Movie S5, Movie S6) that likely contributes to a more efficient repair of damaged lysosomes. Manipulation of PI4P levels using a PI4K2A inhibitor (NC03) or overexpressing Sac1, a phosphatase that metabolizes PI4P at the endolysosomal compartment, corroborated this hypothesis, as both approaches counteracted ARFIP2 depletion and promoted lysosomal damage (Fig. 5g-h, Extended Data Fig. 5g-h).

**Fig. 5.**
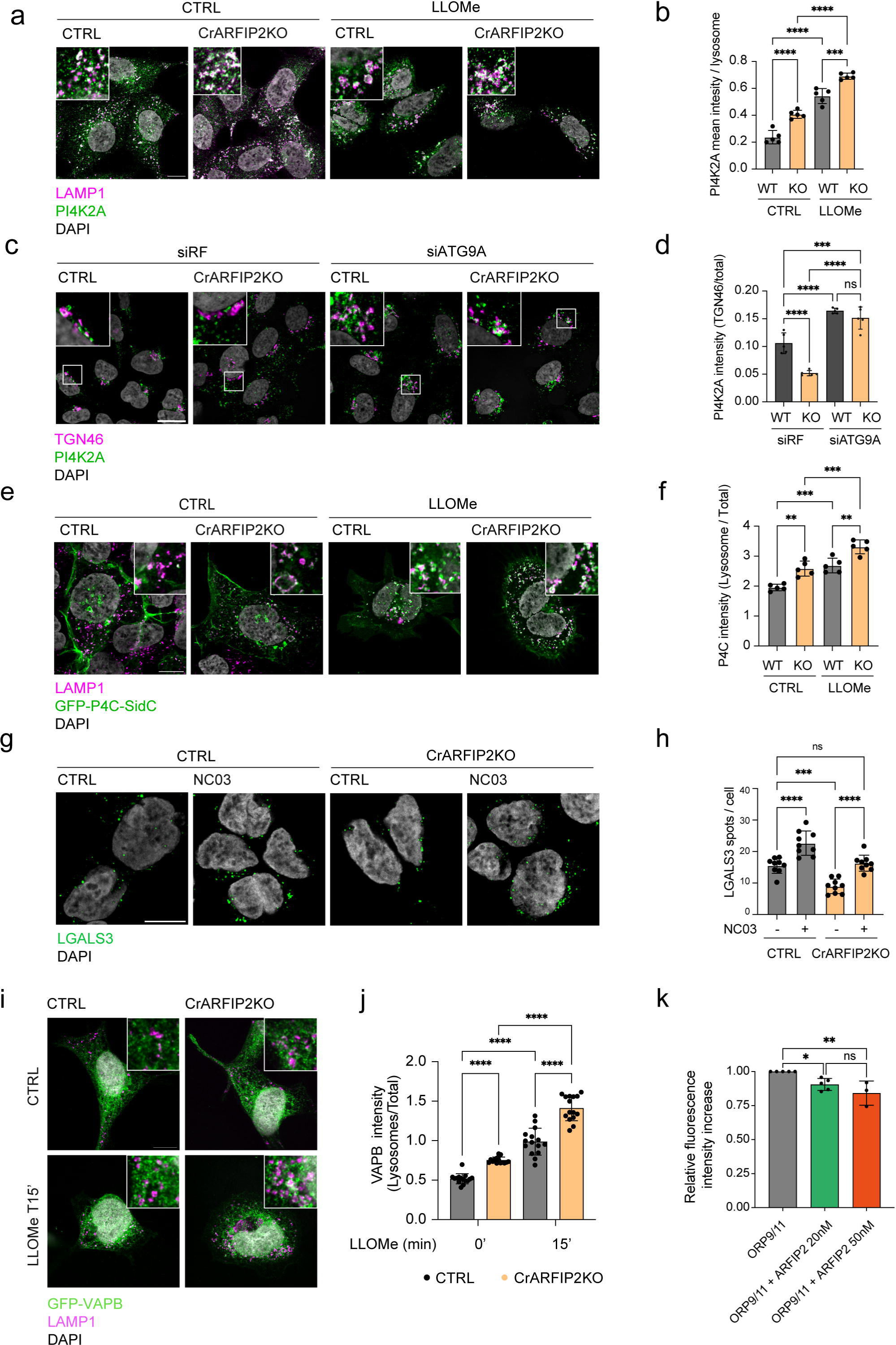
ATG9A and ARFIP2 control PI4K2A lysosomal trafficking and subsequent lysosomal repair. (a) Immunofluorescence of HEK293A CTRL and CrARFIP2KO cells using the indicated antibodies. Scale bar: 10 μm (b) Quantification of PI4K2A in the lysosomal compartment in the indicated cell lines. *n* = 4 independent experiments, **** p < 0.0001. (c) Immunofluorescence of CTRL or CrARFIP2KO cells transfected with siRF or siRNA ATG9A for 72 hours. Scale bar: 10 μm (d) Quantification of PI4K2A in the peri-Golgi area in the indicated cell lines. *n* = 4 independent experiments, *** p < 0.001, **** p < 0.0001. (e) Immunofluorescence of CTRL and CrARFIP2KO cells transfected with GFP-P4C-SidC and treated with 1mM LLOMe for 15 minutes, using the indicated antibodies. Scale bar: 10 μm (f) Quantification of GFP-P4C-SidC intensity in the lysosomal compartment in the indicated cell lines. *n* = 5 independent experiments, ** p < 0.01 *** p<0.001. (g) CTRL or CrARFIP2KO cells were treated or not with PI4K2A inhibitor NC03 25 uM for 1h and subsequently treated with 1mM LLOMe for 15 minutes, with or without NC03. LGALS3 spots were detected by immunofluorescence. Scale bar: 10 μm. (h) Quantification of LGALS3 spots in CTRL cells and cells treated with NC03. *n* = 3 independent experiments, *** p < 0.001, **** p < 0.0001. (i) CTRL and CrARFIP2KO cells were transfected with GFP-VAPB and subsequently treated with 1 mM LLOMe for 15 minutes. Images were acquired using SoRa super-resolution microscope followed by deconvolution. Scale bar: 10 μm. (j) Quantification of VAPB intensity on the lysosomal compartment in CTRL and CrARFIP2KO cells upon LLOMe treatment. *n* = 3 independent experiments, ** p < 0.01, **** p < 0.0001. (k) PS transfer activity of ORP9/11heterodimer in the presence or absence of ARFIP2 at the indicated concentrations using acceptor liposomes enriched with PI4P. *n* = 5 independent experiments.

### ARFIP2 modulated ORPs-mediated lipid transfer for lysosomal repair

PI4K2A-mediated PI4P production is required to establish ER-lysosome contact sites and mediate lysosomal repair via lipid transfer through PI4P binding proteins. Specifically, PI4P is exchanged with phosphatidylserine (PS) or cholesterol by ORP9-10-11 heterodimers and OSBP, respectively. Both ORP9 and OSBP contain an FFAT domain that anchors them to the ER via interaction with VAPA/B. Thus, we investigated whether an increased PI4P production on lysosomes in CrARFIP2KO cells led to a higher number of ER-lysosome contact sites. As expected, immunofluorescence experiments revealed an increased clustering of the VAPB-positive compartment around lysosomes following LLOMe treatment (Fig. 5i-j). Interestingly, VAPB- enrichment on lysosomes was even more pronounced in CrARFIP2KO cells compared to CTRL cells, with or without LLOMe (Fig. 5i-j), suggesting an increased lipid transfer through ER-lysosomes contact sites which should facilitate a more efficient repair.

ARFIP2 AH and BAR domain specifically binds PI4P^23^. Thus, we hypothesized that ARFIP2 might reduce the accessibility of PI4P for ORPs-mediated lipid transfer by binding PI4K2A-mediated PI4P enriched membranes. *In vitro* lipid transport experiments using a fluorescence resonance energy transfer (FRET)-based assay demonstrated that ORP9/11 heterodimer transports PI4P (Extended Data Fig. 5i-j) and such transport is increased when the donor lipid compartment is enriched in PI4P (Extended Data Fig.5j-k). Interestingly, the addition of ARFIP2 significantly inhibits ORP9/11 lipid transport only when the donor liposomes were enriched in PI4P, likely through direct protein-lipid interaction (Fig. 5k, Extended Data Fig. 5j-k).

In summary, our findings suggest that upon damage ATG9A vesicles relocate to lysosomes to deliver PI4K2A, facilitating the production of PI4P required for ER-lysosome contact site formation and lipid exchange for lysosomal repair.

ARFIP2-dependent membrane repair pathway is required for *M. tuberculosis* and *Salmonella* restriction.

In the context of infection, some pathogens use the rupture of membranes to infect the cytosol where they can proliferate. As ARFIP2 emerged as a novel modulator of the PITT pathway, we tested whether it may also have implications for the defence mechanism against pathogens.

To evaluate if the ARFIP2-dependent membrane repair pathway contributes to the restriction of intracellular pathogens, we used primary human monocyte-derived macrophages (HMDM) infected with *Mycobacterium tuberculosis* (Mtb). We confirmed that upon LLOMe treatment, ATG9A was recruited on the lysosomes in human macrophages (Extended Data Fig. 6a), revealing a conserved mechanism for ATG9A. We nucleofected HMDM with Cas9 protein and single guide RNA (sgRNA) targeting ARFIP2^57^ to obtain an ARFIP2 KO pool (ARFIP2^nf^) (Extended Data Fig. 6b-d). Strikingly, after Mtb WT infection there was enhanced bacterial restriction in ARFIP2^nf^ compared to CTRL cells (Fig. 6a-b). In agreement with these results, a reduced number of LGALS3 spots was observed in ARFIP2^nf^ compared to CTRL macrophages upon LLOMe treatment (Fig. 6c-d). These data indicate that loss of ARFIP2 contributes to the restriction of Mtb infection, likely through an ATG9A- mediated membrane repair mechanism.

**Fig. 6.**
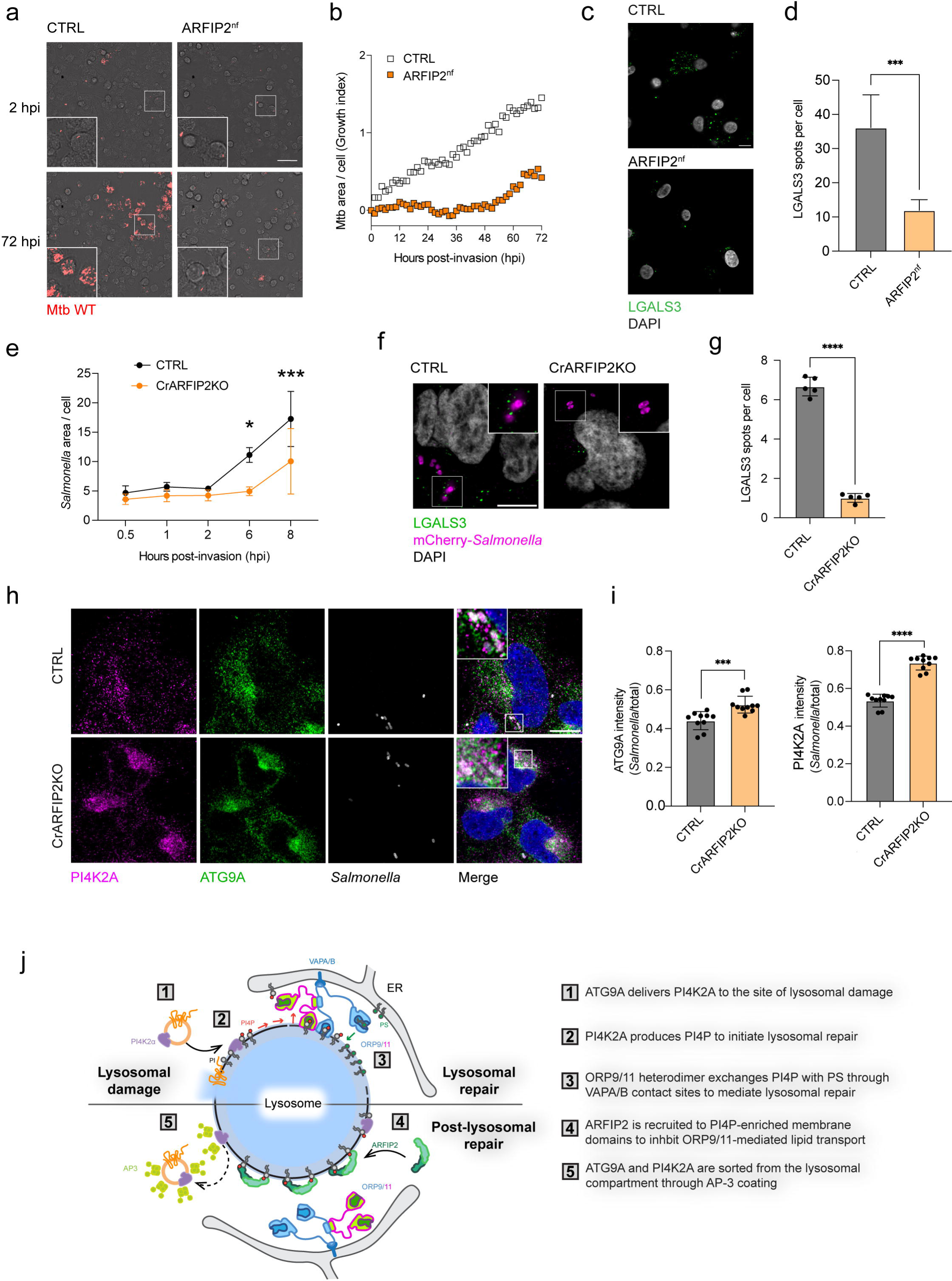
ARFIP2 loss restricts Mtb and *Salmonella* infection. (a) Snapshot of live **H**MDM (CTRL and ARFIP2^nf^) cells infected with Mtb WT at 2[h and 72 h post-invasion. Bright-field and Mtb-E2- Crimson (red). Scale bar: 50[µm. (b) High-content quantitative analysis of live Mtb WT replication in HMDM CTRL and ARFIP2^nf^ cells. Mtb area (dot plot) was calculated as fold change, relative to Mtb uptake at time 0[h post-invasion. *n* = 3 independent experiments. (c) CTRL and ARFIP2^nf^ HMDM cells were treated with 1 mM LLOMe for 15 minutes and LGALS3 spots were detected by immunofluorescence need labels. Scale bar: 10 μm. (d) Quantification of LGALS3 spots in CTRL and ARFIP2^nf^ HMDM cells. *n* = 3 independent experiments, *** p < 0.001 (e) CTRL and CrARFIP2KO cells were infected with mCherry-*Salmonella* for 10 minutes. *Salmonella* growth was monitored as area *Salmonella* area per cell at the time points indicated. *n* = 3 independent experiments, * p < 0.05, *** p < 0.001. (f) CTRL and CrARFIP2KO cells were infected with mCherry-*Salmonella* for 10 minutes. After 30 minutes, immunofluorescence was performed to detect lysosomal damage. Scale bar: 10 μm (g) Quantification of LGALS3 spots in CTRL and CrARFIP2KO after 30 minutes post-invasion with *Salmonella*. *n* = 3 independent experiments, **** p < 0.0001. (h) CTRL and CrARFIP2KO cells were infected with mCherry-*Salmonella* for 10 minutes. After 30 minutes, immunofluorescence was performed to detect ATG9A and PI4K2A recruitment to the pathogen. Scale bar: 10 μm (i) Quantification of ATG9A or PI4K2A intensity on *Salmonella* bacilli. *n* = 3 independent experiments, *** p < 0.001, **** p < 0.0001. (j) Schematic model of ARFIP2-ATG9A-PI4K2A interplay in repairing damaged lysosomes. The different steps of the repair mechanism are described as reported (right).

We also investigated the effects of ARFIP2 loss in *Salmonella* infection. Despite both CTRL and CrARFIP2KO cells showing the same number of cells infected by *Salmonella* infected cells (Extended Data Fig. 6e), pathogen replication was more restricted in CrARFIP2KO cells compared to CTRL cells (Fig. 6e, Extended Data Fig. 6f). Further, similarly to LLOMe treatment, CrARFIP2KO cells showed a reduced number of LGALS3 spots during *Salmonella* infection (Fig. 6f-g), together with an increased association of ATG9A and PI4K2A with bacteria (Fig. 6h-i). Additionally, mCherry-*Salmonella* was engulfed in LAMP1-positive membranes in CrARFIP2KO cells (Extended Data Fig. 6g), confirming that the pathogen was restricted in the endolysosomal compartment upon ARFIP2 loss. Overall, these data support a role for ARFIP2 in regulating the retrieval of ATG9A and PI4K2A from the site of pathogen infection after membrane repair.

## DISCUSSION

Lipid metabolism has gained a central role in the process of lysosomal repair upon damage where lipids are needed to expand and seal the damaged membranes. Lipid metabolizing enzymes together with the formation of membrane contact sites between lysosomes and other membrane compartments are now acknowledged as key factors to counteract LMP. Our work elucidates a new mechanism by which ATG9A vesicles transiently interact and potentially fuse with lysosomes after LMP, delivering PI4K2A to the site of lysosomal damage to produce PI4P and initiate the PITT pathway for lysosomal repair. Loss of ATG9A blocks PI4K2A in the peri-Golgi compartment preventing its function at the lysosomes and increasing lysosomal damage. Previous studies have shown that ATG9A regulates the trafficking of another PI-metabolizing enzyme, PI4KIIIB ^16^,that has also been reported to localize at the lysosomes ^58^. However, while the pool of PI4P produced by PI4K2A has been described to be crucial for lysosomal repair ^6^, PI4P produced by PI4KIIIB has been shown to be dispensable for lysosomal damage and repair and more relevant for lysosomal reformation coupled with ARF-1 ^58^.

How ATG9A is mobilized to the lysosomes upon LMP is still unresolved. Our data ruled out autophagy initiation or ATG8s lipidation by CASM as stimuli that trigger ATG9A trafficking to the lysosomal compartment: indeed, WIPI2 puncta are not detected in the early steps of LMP when ATG9A is already at the lysosome. Furthermore, ATG9A is redirected to the lysosomal compartment also in HeLa Hexa KO cells upon sterile damage, where all ATG8s are absent supporting an independent mechanism for ATG9A lysosomal localization. While the trigger for ATG9A mobilization is still unclear, we have uncovered a new mechanism by which ATG9A is retrieved from the lysosome which involves the coordinated action of ARFIP2 and AP-3.

In addition, we describe a new role for ARFIP2 during lysosomal damage and repair. Loss of ARFIP2 induces increased lysosomal repair by affecting ATG9A trafficking and leading to an accumulation of ATG9A, and in turn of PI4K2A, in the lysosomal compartment. Indeed, these effects are reverted by the loss of ATG9A. As ARFIP2 is a PI4P interacting protein and requires PI4P binding via the AH to exert its functions in lysosomal repair, we propose a model where ARFIP2 is recruited through PI4P to the damaged lysosomes to promote the retrieval of ATG9A back to the Golgi when the repair commences. It is still uncertain whether ATG9A might function as a scramblase at the lysosomes, a hypothesis that future studies should address. Indeed, ATG9A may potentially facilitate, by means of its scramblase activity, the accessibility of lipids that are crucial for lysosomal repair by ORPs-mediated lipid transfer through ER-lysosomes contact sites. Alternatively, ATG9A could bind ATG2 modulating its lipid transfer activity to fine-tune the flux of lipids needed to repair the lysosomal membranes after damage. Both of these mechanisms require further work.

Here, we propose a model where, after LMP, the production of PI4P by PI4K2A reaches a critical concentration, leading to the recruitment of ARFIP2 to PI4P-enriched lysosomal domains through its AH-BAR domain, thereby inhibiting PI4P lipid transfer mediated by ORP9-10-11 heterodimers and OSBP (Fig. 6j). Concurrently, through its N-terminal domain, ARFIP2 along with PI4K2A, interact with AP-3 to facilitate the coat assembly on PI4P-enriched regions, where ATG9A-positive vesicles will be formed and retrieved to the Golgi. Indeed, using different biochemical approaches, we have validated AP-3 as a new adaptor complex that decorates ATG9A vesicles and supports the sorting from the lysosomal compartment. AP-1, AP-2 and AP-4 have also been described as regulators of ATG9A vesicle trafficking in other membrane compartments. ATG9A contains both a tyrosine motif and a dileucine motif that enables its interaction with AP-1, AP-2 and AP-4. However, we could not detect a direct interaction between AP-3 and ATG9A, suggesting that other proteins, such as PI4K2A and ARFIP2, might be mediating the assembly of the coat on ATG9A-positive membrane regions. Since, in all probability different adaptor complexes cannot coat the same vesicle simultaneously, we suggest that each adaptor complex identifies a distinct pool of ATG9A vesicles which is devoted to a specific compartment-specific functions.

Finally, we have translated the importance of this pathway in the physiological context of pathogen infection. During infection, pathogens are directed to phagosomes to facilitate their elimination through endolysosomal degradation^59^. However, several intracellular bacteria can damage the membrane of phagosomes and access the cytosol^60^. Maintaining the functionality of the endolysosomal compartment is crucial to control the infection. In this scenario, the canonical repair mechanism mediated by ESCRT prevents lysosome rupture during pathogen infection^61^. Recently, PI4P and OSBP have been described to accumulate on *Mycobacterium*-containing vacuole (MCV) establishing ER-contact sites to restore MCV integrity by an ER-dependent repair mechanism^62^. Our data demonstrate that loss of ARFIP2 increased the restriction of Salmonella infection in HEK293A cells and Mtb in human macrophages, revealing the broad significance of this regulatory mechanism of lysosomal repair regulatory mechanism across various cell types and different pathogens. These data, along with the mobilization of ATG9A upon LMP induction in different cell lines, warrant further investigation in other biological contexts where lysosomal damage and repair plays a critical role, such as neurodegenerative diseases or cancer.

## Materials and Methods

### Cell culture

HEK293A human embryonic kidney cells (CRL-1573) and *Mocha* fibroblasts (CRL-2709) were obtained from American Type Culture Collection (ATCC). Mouse embryonic fibroblasts (MEF) and SH-SY5Y cells were obtained from Cell Services at The Francis Crick Institute.

Cells were grown in DMEM high glucose (Sigma, D6429) supplemented with 10% heat-inactivated fetal bovine serum (FBS) (Gibco, 10270-106). Cells were incubated at 37 °C, 10% CO_2_ and 90-95% of relative humidity. Specific experimental conditions are indicated in figure legends. The ARFIP2 KO cell line was generated by CRISPR/Cas9-mediated genome engineering as previously described^16^. The ATG9A KO cell line was generated by CRISPR/Cas9-mediated genome engineering as previously described^33^. All chemicals and reagents used in this study are reported with the working concentrations in Table S3.

SH-SY5Y 14-day differentiation protocol was performed as previously described^63, 64^. Briefly, on day 1 and 4, the media was switched to DMEM high glucose supplemented with 2.5% heat-inactivated FBS and 10 μM Retinoic Acid (Merck, R2625). On day 7, cell culture plates were coated with laminin (Corning, 10152421) at 2 μg/cm^2^. On day 8 and 11, the media was switched to a neuron differentiation media consisting of Neurobasal Medium (Thermo Fisher Scientific, 21103049) supplemented with 1x B-27 Supplement (Thermo Fisher Scientific, 17504044), 50 ng/mL Human Recombinant brain-derived neurotrophic factor (BDNF) (Stem Cell Technologies, 78005), 0.2 mM dibutyryl cyclic AMP (db-cAMP) (Merck, 28745), 20 mM Potassium chloride (Sigma-Aldrich, P9531), 2 mM L-Glutamine (Sigma-Aldrich, G7513), and 10 μM Retinoic Acid. On day 14, cells were processed for the specific experiments.

### Preparation and culture of HMDMs

Human monocytes were prepared from leucocyte cones (NC24) supplied by the NHS Blood and Transplant service^60^. White blood cells were isolated by centrifugation on Ficoll-Paque Premium (17-5442-03; GE Healthcare) for 60 min at 300 × *g*. Mononuclear cells were collected and washed twice with MACS rinsing solution (130-091-222; Miltenyi) to remove platelets and red blood cells. The remaining samples were incubated with 10 ml RBC lysing buffer (R7757; Sigma-Aldrich) per pellet for 10 min at room temperature. Cells were washed with rinsing buffer and were resuspended in 80 µl MACS rinsing solution supplemented with 1% BSA (130-091-376; MACS/BSA; Miltenyi) and 20 µl anti-CD14 magnetic beads (130-050-201; Miltenyi) per 10^8^ cells. After 20 min on ice, cells were washed in MACS/BSA solution and resuspended at a concentration of 10^8^[cells/500 µl in MACS/BSA solution and further passed through an LS column (130-042-401; Miltenyi) in the field of a QuadroMACS separator magnet (130-090-976; Miltenyi). The LS column was washed three times with MACS/BSA solution, then CD14 positive cells were eluted, centrifuged, and resuspended in complete RPMI 1640 with GlutaMAX and Hepes (72400-02; Gibco) and 10% fetal bovine serum (FBS; F7524; Sigma-Aldrich).

### Nucleofection of HMDM

Human monocytes were washed twice with PBS and electroporated in the appropriate primary nucleofection solution (Cat. No. VPA-1007; Amaxa Human Monocyte Nucleofector Kit) using the Lonza 2b Nucleofector (AAB-1001; Nucleofector 2b Device). 5 × 10^6^ of human monocytes were used per reaction and resuspended in 100 µl of primary nucleofection solution containing 4 µg of S.p. Cas9 (IDT) mixed with a total of 12 µg of targeting synthetic chemically modified sgRNAs (Synthego; Table S4). Human monocytes were then nucleofected with the sgRNA pool and the Cas9-RNP mix using the Y001 program. Nucleofected cells were cultured in prewarmed RPMI 1640 supplemented with GlutaMAX, Hepes, and 10% FBS in a 6-well plate. 2 h after nucleofection, 100 ng/ml hM-CSF was added to the cells. Dishes were incubated in a humidified 37°C incubator with 5% CO_2_. After 3 d, an equal volume of fresh complete media including 100 ng/ml hM-CSF was added. 6 d after the initial isolation, differentiated macrophages were detached in 0.5 mM EDTA in ice-cold PBS using cell scrapers (83.1830; Sarsted), pelleted by centrifugation, and resuspended in RPMI medium containing 10% FBS^65^.

### Plasmids and siRNA transfection

DNA transfection was performed following manufacturer’s instructions using Lipofectamine 2000 (Invitrogen, 11668-019) in 1:5 Opti-MEM:DMEM medium (Gibco, 31985-047). Plasmids generated for this study are available upon request: pcDNA3.1-mCherry-3xFLAG-His-TEV-ATG9A, pcDNA3.1-myc-TurboID, pcDNA3.1-myc-ATG9A-TurboID. pBMN-AP-3D1 was kindly provided by Dr A. Peden^44^. pGEX6p-1 GST-ARFIP2, pDEST-GFP-ARFIP2 and GFP-BAR were a gift from K. Nakayama ^22^ (Kyoto University, Kyoto, Japan) and GFP-ARFIPs Chimera was generated from pDEST-GFP-ARFIP2. PI4K2A plasmids were a gift from Shane Minogue (UCL, Institute for Liver and Digestive Health, London, United Kingdom). pEGFP-N1 TGN46-GFP was a gift from V Ponnambalam (University of Leeds, United Kingdom). pcDNA3.1-mCherry-3×FLAG-6×His-TEV- ATG9A was purchased from Genescript. GFP-Sac1 and GFP-P4MX2 were kindly gifted from G. Hammond ^66^ (University of Pittsburgh, Pittsburgh, PA). EGFP-P4C-SidC plasmid was a gift from Michael Marks (University of Pennsylvania, Philadelphia, USA) and Tamas Balla (NIH, Bethesda, USA). GFP-VAPB plasmid was a gift from Tim Levine (UCL Institute of Ophthalmology, London,

United Kingdom). pC3 ORP9 and ORP11 plasmids were kindly gifted by JX Tan ^6^ (Aging Institute, University of Pittsburgh School of Medicine and University of Pittsburgh Medical Center, Pittsburgh, PA, USA).

### Mycobacterial strains and culture conditions

Mtb H37Rv (Mtb WT) was kindly provided by Prof. Douglas Young (The Francis Crick Institute, London, UK). Fluorescent Mtb strains were generated as previously reported^67^. E2Crimson Mtb was generated by transformation with pTEC19 (30178; Addgene, deposited by Prof. Lalita Ramakrishnan). The strain was verified by sequencing and tested for phthiocerol dimycocerosate positivity by thin-layer chromatography of lipid extracts from Mtb cultures. Mtb strain was cultured in Middlebrook 7H9 (M0178; Sigma-Aldrich) supplemented with 0.2% glycerol (G/0650/17; Fisher Chemical), 0.05% Tween-80 (P1754; Sigma-Aldrich), and 10% ADC (212352; BD Biosciences).

### Macrophage infection with Mtb

The day before infection, HMDM were seeded at a density of 60,000 cells per well of a 96-well plate. Mid-logarithmic phase bacterial cultures (OD_600_ 0.5-1.0) were centrifuged at 2,000 × *g* for 5 min and washed twice in PBS. Pellets were then shaken vigorously for 1 min with 2.5–3.5 mm glass beads (332124G; VWR) and bacteria were resuspended in 10 ml macrophage culture media before being centrifuged at 300 × *g* for 5 min to remove large clumps. The top 7 ml of bacterial suspension was taken, OD_600_ recorded and diluted appropriately for infection. The inoculum was added at the correct MOI, assuming OD_600_ of 1 is 1 × 10^8^ bacteria/ml. Infections were carried out in a volume of 50 µl in a 96-well plate, 300 µl in a 24-well plate, or 500 µl in a 12-well plate. After 2 h of uptake, extracellular bacteria were removed with two washes in PBS and macrophages were incubated at 37°C and 5% CO_2_ for the required time points in macrophage media. An MOI of 1 was used for replication experiments.

### Long-term live-cell imaging of Mtb replication and HMDM

For live-cell imaging, 60,000 macrophages were seeded per well on an olefin-bottomed 96-well plate (6055302; Perkin Elmer). Cells were infected with Mtb at an MOI of 1 for 2 h. After infection, cells were washed with PBS and replaced with a macrophage media. Imaging was performed using the OPERA Phenix microscope with a 40× 1.1 NA water-immersion objective with a 10% overlap between adjacent fields. Five planes with 1 µm distance of more than 20 fields of view were monitored in time and snapshots were taken every 1.5 h for 72 h. For imaging on the Opera Phenix, Brightfield was detected using λex = transmission/λem = 650–760 nm, and E2-Crimson bacteria was detected using λex = 640 nm/λem = 650–760 nm using a 16-bit scMOS camera. For assessing bacterial replication, analyses were performed with Harmony software where maximum projection of individual z-planes with an approximate distance of 1 µm was used. To perform cellular segmentation “Find texture regions,” building blocks were trained in Brightfield channel to segment cellular areas. Following the segmentation of cellular area Find spots, building blocks were used to segment Mtb. To determine the bacteria area over time, the spot area was summed for each time point. Mtb replication as growth index was calculated by the formula: (sum of intracellular Mtb area for the time point − sum of intracellular Mtb area t0[h) / (sum of intracellular Mtb area t0[h).

### *Salmonella* infection

*Salmonella enterica* serovar Typhimurium, strain 14028swere used for all cell culture studies. mCherry-expressing Salmonella was grown with 50 μg/mL carbenicillin. 50 μg/mL kanamycin was added for the culture of bacteria. For SPI-1 induced infection of HEK293A cells, bacteria were grown overnight in Luria broth (LB) and sub-cultured (1:33) in fresh LB for 3.5 h prior to infection at 37 °C. Cells seeded in 24-well were infected with 10 μL of *Salmonella* subculture, for 10 min at 37 °C. After two PBS washes, cells were incubated in 100 μg/mL gentamycin for 1-2 h and 20 μg/mL gentamycin thereafter.

### Proximity labelling using TurboID and streptavidin pull-down

Stable myc-TurboID and myc-ATG9A-TurboID HEK293A cells were cultures in 15 cm Petri dishes until they reached 80% confluency. Cells were treated with 50 µM Biotin in complete medium to allow biotinylation at different time points (1 hr for MS experiments and 15 minutes for Western Blot validation). After Biotin treatment, cells were washed 3 times with ice-cold PBS and centrifuged to generate cell pellets that were frozen. For Western Blot analysis, lysates were lysed in 700 µL TNTE lysis buffer [20 mM Tris-HCl pH 7.4, 5 mM EDTA, 150 mM NaCl, 0.5% Triton- X100] supplemented with EDTA-free protease (Roche, cOmplete EDTA-free, 05056489001) and phosphatase inhibitors (Roche, Phostop EASYpack, 04 906 837 001). Cell lysates were centrifuged at 13000 x g for 10 min at 4 °C. Lysates were pre-cleared with 15 µL of empty agarose beads in a wheel for 1h at 4 °C. Protein concentration was determined by Bradford (Bio-Rad, Bio-Rad Protein Assay Dye Reagent Concentrate, 5000006) and protein amounts were normalised among samples. Inputs 2% were taken for Western Blot analysis. Pre-cleared lysates were subjected to Streptavidin pull-down by incubating samples with 30 µL washed beads in a wheel for 2 h at 4 °C supplemented with 1% SDS. After pull-down, beads were washed three times (10 minutes each) at 4 °C and eluted at 65 °C for 15 minutes in elution buffer (Laemli buffer + 3 mM Biotin). Samples were further processed for either Western Blot or label-free MS as described in the corresponding sections.

### GFP-trap immunoprecipitation

Cells were lysed in ice-cold TNTE buffer (20 mM Tris, pH 7.4, 150 mM NaCl, 1%w/v TritonX-100, 5 mM EDTA) containing EDTA-free Complete Protease Inhibitor cocktail (Roche). Lysates were cleared by centrifugation at 21000 x g and precleared with binding control agarose beads (ChromoTek) for 1 hour at 4 °C. GFP-tagged proteins were immunoprecipitated using GFP-TRAP beads (ChromoTek) for 2h at 4 °C. After immunoprecipitation, GFP-beads were washed three times with ice-cold TNTE buffer and resuspended in 2× Laemmli sample buffer. Samples were further processed for either Western Blot or label-free MS as described in the corresponding sections.

### Cell lysis and Western Blot

For cell lysis, cells were washed three times with ice-cold PBS, scraped and lysed on Lysis Buffer [20 mM Tris-HCl pH 7.4, 5 mM EDTA, 150 mM NaCl, 0.5% Triton-X100] supplemented with EDTA-free protease (Roche, cOmplete EDTA-free, 05056489001) and phosphatase inhibitors (Roche, Phostop EASYpack, 04 906 837 001). Cell lysates were centrifuged at 13000 x g for 10 min at 4 °C. Protein concentration was analyzed using Pierce BCA Protein Assay kit (Thermo Scientific, 23227) following manufacturer’s instructions. Equal amounts of protein lysates were resuspended in Laemnli SDS-sample buffer and incubated at 65 °C for 10 min. Proteins were resolved on NuPage 4- 12% Bis-Tris Gel (Invitrogen, NP0336) using MES SDS running buffer (Invitrogen, NP0002) and transferred to Immobilon-P transfer membranes (Millipore, IPVH00010). Membranes were blocked with 5% non-fat dry milk (BioRad, 1706404) in Phosphate-buffered saline containing 0.1% Tween- 20 (PBS-T) for 1 h at room temperature. Incubation of primary antibodies was performed overnight at 4 °C in 5% non-fat dry milk or 3.5% BSA (Roche, 10735086001). After three washes in PBS-T, membranes were incubated for 1 h at room temperature with secondary antibodies (1:5000) diluted in 5% non-fat milk. Upon incubation, membranes were washed three times with PBS-T and protein detection was performed by using Immobilon Classico Western HRP Substrate (Millipore, WBLUC0500) or Immobilon Crescendo Western HRP Substrate (Millipore, WBLUR0500). Blots were scanned with Amersham ImageQuant 800 (Cytiva). Densitometry analysis of Western Blots was performed using FIJI (https://fiji.sc/). All the antibodies used in this study are reported with the working concentrations in Table S3.

### Mass spectrometry LC-MS/MS and MS data processing and analysis

For the Proximity labelling assay using TurboID and streptavidin pull-down MS experiments were performed by DDA analysis on Orbitrap Fusion Lumos. Peptides were analysed using an Evosep One LC system (EvoSep Biosystems) directly coupled to an Orbitrap Fusion Lumos tribrid mass spectrometer (Thermo Scientific). Reverse phase separations were performed at a flow rate of 500 nL/min on an EV1064 ENDURANCE analytical column (100 μm × 8 cm, 3.0 μm particle size; Evosep Biosystems) using the vendor’s predefined 30 samples per day gradient method. The Orbitrap was operated in ‘TopS’ Data Dependent Acquisition mode with precursor ion spectra acquired at 120k resolution in the Orbitrap detector and MS/MS spectra at 32% HCD collision energy in in the ion trap. Automatic Gain Control was set to Auto for MS1 and MS2. Maximum injection times were set to ‘Standard’ (MS1) and ‘Dynamic’ (MS2). Dynamic exclusion was set to 20s. For the DDA Data Processing and Analysis MaxQuant (version 1.6.12.0) was used for data processing. The data was searched against the *Homo Sapien* UniProt reference proteome. A decoy database containing reverse sequences was used to estimate false discovery rates and set the false discovery rate at 1%. Default MaxQuant parameters were used with the following adjustments: Label-free quantification was selected along with iBAQ values, with Normalization type “Classic” selected. MaxQuant output files were imported into Perseus (version 1.4.0.2) and the LFQ intensities and iBAQ values were used for all subsequent analysis. Missing values were imputed from a normal distribution.

For the GFP-trap immunoprecipitation experiments, MS experiments were performed by Data-independent acquisition (DIA) analysis on Fusion Lumos Orbitrap. Peptides were analysed using an Evosep One LC system (EvoSep Biosystems) directly coupled to an Orbitrap Fusion Lumos tribrid mass spectrometer (Thermo Scientific). Reverse phase separations were performed at a flow rate of 500 nL/min on an EV1064 ENDURANCE analytical column (100 μm × 8 cm, 3.0 μm particle size; Evosep Biosystems) using the vendor’s predefined 30 samples per day gradient method. Lumos instrument settings were as follows: MS1 data acquired in the Orbitrap with a resolution of 120 k, max injection time of 20 ms, AGC target of 1e6, in positive ion mode, in profile mode, over the mass range of 393 to 907 *m/z*. DIA segments over this mass range (20 *m/z* wide/1 Da overlap/27 in total) were acquired in the Orbitrap following fragmentation in the HCD cell (32%), with 30 k resolution over the mass range 200 to 2,000 *m/z* and with a max injection time of 54 ms and AGC target of 1e6. For the DIA-MS Data Processing and Analysis, the data were searched using Direct DIA data analysis on Spectronaut v.14 (Biognosys AG) using default settings, then run-wise imputation (*Q*- value percentile = 30%) were applied to the dataset. A two-sample *t*-test was carried out in Spectronaut software, then filters were applied to the data (*q* ≤ 0.05, avg log_2_-fold change ≥ 0.58, no. unique total peptides ≥ 2).

### Immunofluorescence and confocal microscopy

Cells were grown on poly-D-lysine treated coverslips at a 70% confluency the day of the experiments. After the treatments, cells were fixed with 4% paraformaldehyde in phosphate-buffered saline (PBS) supplemented with 0.1 mM CaCl_2_, 0.1 mM MgCl_2_ for 10 min. Cells were washed three times with PBS before adding 50 mM NH_4_Cl for 10 min at room temperature and then permeabilized with 50 μg/mL digitonin (Merck Millipore; D141) for 5 min at room temperature. Coverslips were then washed three times with PBS before the addition of the blocking solution (5% BSA in PBS) for 30 min at room temperature. Coverslips were incubated upside down with primary antibody diluted in 1% BSA/PBS for 1 h at room temperature, then washed three times with PBS and incubated upside down with secondary antibody in 1% BSA/PBS for 1 h. Finally, coverslips were washed three times in PBS and once with demineralized water before mounting them on glass microscope slides with 10 μL Mowiol mounting solution per coverslip. Fluorescence images were acquired using a Zeiss LSM 880 Airyscan confocal microscope with Plan-Apochromat 63x/1.4 Oil DIC M27 objective lens. Zeiss ZEN imaging software was used for the acquisition. Antibodies used in this study with the working concentrations are listed in Table S3. For SoRa imaging, images were acquired on a Nikon SoRa spinning disk, using a 60X/1.4 Oil immersion objective and SoRa magnification disk to get a final pixel size of 27 nm x 27 nm. The different channels were acquired exciting with the laser lines 405 nm, 488 nm, and 561 nm, and detecting with selective bandpass filters for DAPI (447/60 nm), Alexa Fluor 488 (525/50 nm), and Alexa Fluor 555 (600/52 nm).

### Lysotracker uptake and recovery

Twenty thousand cells were seeded into 96-well plates (Greiner Bio One Ltd 655090). Cells were loaded with the nuclear dye Hoechst 33342 (Thermo Fisher Scientific) at a dilution of 1:10,000 and 25[nM LysoTracker DND-99 (Thermo Fisher Scientific; L7528) for 45[minutes. The cells were imaged every 1 min at 37[°C, 5% CO2 using an Opera Phenix microscope (PerkinElmer). First, a baseline was established by imaging 3 time points corresponding to 20 minutes, followed by the addition of LLOMe to a final concentration of 1[mM. After 15 minutes, the cells were washed 3 times with DMEM and the medium was replaced with DMEM containing 25[nM LysoTracker, and lysosomal recovery was followed for 80 min. Image acquisition and analysis were performed as indicated below (see ‘Live Imaging’).

### Live imaging

Cells transfected with mCherry-3xFLAG-ATG9A or mCherry-3xFLAG-ATG9A/GFP-TGN46 were seeded on glass-bottom microwell dishes (MatTek Corp.; P35G-1.5-14-C) to reach a 70% confluency the day of the experiment. For the LysoTracker and dextran uptake experiments, 30 min before image acquisition, transfected cells were loaded with Lysotracker Blue DND-26 (Thermo Fisher Scientific; L7525) and dextran Alexa-fluor 488 10,000 MW (Invitrogen; D22910) diluted in full medium at the appropriate concentration according to manufacturer’s instructions. Cells were imaged using a Zeiss LSM 880 Airyscan confocal microscope with Plan-Apochromat 63x/1.4 Oil DIC M27 objective lens. Live imaging was performed using Zeiss ZEN imaging software. After the acquisition, movies were processed using an Airyscan processing tool on the ZEN software provided by Zeiss. For the Lysotracker uptake assay, the microscope was pre-warmed at 37 degrees and supplied with 5% CO2, prior to imaging. Cells were imaged with 63x/NA (1.15) water-immersion lens. 4 Z-stacks with a step size of 1µm were imaged were acquired using excitation lasers at 375, and 568 nm, and emission filters at 435-480, and 570-630 nm, respectively. Cell segmentation and quantification analysis were performed using Harmony software 5.0.

### Image analysis

Image analysis has been performed using the open-source FIJI (http://fiji.sc) ^68^ and the pipelines for the different quantified phenotypes have been designed as follows: (i) ATG9A or PI4K2A localization in the Golgi. ATG9A image was used to threshold and generate a binary image of total fluorescence. GM130 was used to generate a mask of the Golgi region. The ratio between Golgi ATG9A and total ATG9A was calculated as previously described^46^, (ii) ATG9A intracellular dispersal. In cases where a Golgi marker was not used, a mask of the nuclei was used to create a distance map corresponding to the segmented cells. ATG9A intensity was calculated according to the distance map to estimate the dispersal from the nuclei upon starvation, (iii) ATG9A co-occurrence with AP-3 complex. A line was created using the Line Tool to select a region and an intensity plot was generated for the same segment in the AP-3 channel and the ATG9A channel. The analysis was focused on the region where ATG9A is accumulated, (iv) Vesicle mean velocity was calculated by using the TrackMate2 plugin in FIJI following the indications of the creator (ref), (v) ATG9A/PI4K2A/VAPB localization in the lysosomal compartment. ATG9A image was used to threshold and generate a binary image of total fluorescence. LAMP1 was used to generate a mask of the lysosomes. The ratio between Lysosomal ATG9A and total ATG9A was calculated, (vi) LGAL3 spot counting. LGAL3 channel was used to create a binary image that reflected the positive signal by adjusting the threshold. The “analyze particles” plugin in FIJI was used to determine the number.

Nuclei counting was used to normalize the number of spots per cell, (vii) Salmonella/Mtb area per cell. Salmonella or Mtb were used to mask and create a binary image to calculate their respective total area that was then normalized on the number of nuclei per image (viii) ATG9A/PI4K2A localization in Salmonella. Salmonella was used to mask and create a binary image to calculate the ratio of ATG9A or PI4K2A on the particle versus total ATG9A or PI4KA.

### Immunoisolation of ATG9A-positive membranes

ATG9A-positive membranes were isolated by adapting the protocol established in our laboratory^16^. Briefly, cells were washed in ice-cold PBS and harvested by centrifugation at 200 x g at 4 °C. Pellets were resuspended using ice-cold isotonic buffer (20 mM HEPES, pH 7.4, 250 mM sucrose, and 1 mM EDTA) supplemented with EDTA-free complete protease and phosphatase inhibitors. The resuspended pellet was passed through a 27G needle 15 times for homogenization before clarification by centrifugation at 3,000 x g at 4 °C. Supernatants were incubated overnight at 4 °C with hamster anti-ATG9A or hamster IgM CTRL coupled with protein A Dynabeads (Invitrogen; 10002D). The ATG9A-positive membranes were washed three times with isotonic buffer at 4 °C and resuspended in 2× Laemmli sample buffer before being resolved by SDS-PAGE and Western blotting.<u>9</u>

### Differential centrifugation for cell fractionation

Cells were seeded in two 15-cm dishes per condition to reach 80% confluency the day of the experiment. Cell monolayers were washed twice with ice-cold PBS and scraped in 5 ml PBS followed by centrifugation at 100 x g for 5 minutes to obtain the cell pellet. Cell pellet was resuspended in 700 uL of ice-cold isotonic buffer (20 mM HEPES, pH 7.4, 250 mM sucrose and 1 mM EDTA) supplemented with EDTA-free protease (Roche, cOmplete EDTA-free, 05056489001) and phosphatase inhibitors (Roche, Phostop EASYpack, 04 906 837 001). Cells were mechanically lysed by passing them through a 27G needle attached to a 1 mL syringe on ice. To lyse MEFs, 25 up and down passes were needed. The protocol was adapted from Shoemaker et al, 2019. The post-nuclear supernatant (PNS) was obtained by centrifugation at 1000 x g for 10 minutes at 4 °C to remove nuclei and cell debris. Differential centrifugation protocol using the following speeds were subsequently performed: 3000 x g for 20 minutes, 20000 x g for 30 minutes and 100000 x g for 30 minutes. Fractions 1 and 2 were treated with Benzonase to eliminate any DNA traces and fraction samples were run in SDS-PAGE as described above.

### Protein expression and purification

Arfaptin 2 (pGEX-6P-1-GST) was transformed in Escherichia coli BL21 (DE3) cells. Bacterial were grown at 37 [in LB to an OD600 of 0.6-0.8. Protein expression was induced with 0.5 mM IPTG for 16 hrs at 18[. Cells were harvested by centrifugation and resuspended in ice cold lysis buffer containing 50 mM Tris-HCl pH 7.5, 500 mM NaCl, 0.5 mM TCEP, 0.4 mM AEBSF, and 15 [µg/ml benzamidine. Cells were lysed by freeze-thaw, followed by sonication. Cell lysate was cleared by centrifugation at 25,000g for 30min, at 4[. The protein was absorbed onto 1mL of Glutathione-Sepharose 4B affinity matrix (GE Healthcare) for 2 hrs, and recovered by homemade 3C protease cleavage at 4 [overnight in 50 mM Tris-HCl pH 7.5, 500 mM NaCl, and 0.5 mM TCEP. The eluted protein was further purified by size-exclusion chromatography using Superdex 200 16/60 column (GE Healthcare) equilibrated in buffer containing 25 mM Tris-HCl pH 7.5, 150 mM NaCl, 0.5 mM TCEP and 5% glycerol.

Flag-ORP9/11 heterodimer was expressed and purified from Expi293 cells (ThermoFisher). To transfect cells, cells were grown to a density of 2-3x10^6^/mL on a shaker at 130 rpm, at 37[, supplemented with 8% CO_2_ in Expi293 Expression medium. 1μg of plasmids (ORP9: ORP11, in amount ratio 1:1) per 1x10^6^ cells was mixed with a threefold (w/w) of PEI (polyethyleneimine, linear MW 25000, Polysciences) in Opti-MEM. The transfection mixture was added to the cells, and the cells were grown for another 72 hrs. The transfected cells were pelleted and resuspended in the lysis buffer containing 50 mM Tris-HCl pH 8.0, 500 mM NaCl, 0.5 mM TCEP, 5% glycerol, EDTA-free Complete Protease Inhibitor cocktail (Roche) and 0.1% Triton X-100. Cells were disrupted by three times freeze-thaw cycles in liquid nitrogen and water bath. The cell lysate was cleared by centrifugation at 20,000g, for 30min at 4[. The supernatant was incubated with anti-DYKDDDDK G1 affinity resin (Genscript) for 4-5 hours at 4[. The resin was washed with buffer containing 50 mM Tris-HCl pH8.0, 500 mM NaCl, 0.5 mM TCEP, 5% glycerol and the proteins were eluted with 5 mg/mL 3xFlag peptide dissolved in water. The eluted protein was further concentrated by 30kDa molecular weight cut-off centrifugal concentrator, snap frozen in liquid nitrogen, stored at -80 [.

### Liposome preparation and Lipid transfer assay

Lipids were mixed at the indicated molar ratio in chloroform, dried to a lipid film under nitrogen gas, and further vacuumed for 2 hrs. The lipid film was rehydrated and resuspended in the assay buffer containing 25 mM Tris-HCl pH 8.0, 150 mM NaCl and 0.5 mM TCEP, with vortexing. The resuspended lipid solution was subjected to 5 cycles of freeze-thaw in liquid nitrogen and water batch. The liposome solution was extruded 10 times through 0.2 μm membrane, followed by at least 20 times through 0.1 μm membrane via a Mini-Extruder (Avanti Polar Lipid). The final concentration of liposome was 1 mM. The liposomes had an average diameter of 100 nm, measured by Zetasizer Nano ZS (Malvern Instruments).

Lipid transfer assays were performed at 25 [with at least three independent repeats. Donor liposomes contained 66% DOPC, 25% DOPE, 5% DGS-NTA (Ni), 2% NBD-PS and 2% Rh-PE.

Acceptor liposomes contained 75% DOPC and 25% DOPE, or 70% DOPC, 25% DOPE and 5% brain PI4P. Briefly, the reaction sample (final volume: 80 μL) containing 100 nM ORP9/11 (heterodimer), 20 nM or 50 nM ARFIP2 was mixed with 100μM donor liposomes and 100μM acceptor liposomes. The reaction mix was then transferred into 10mm pathlength quartz cuvette and the NBD fluorescence (excitation at 468 nm, emission at 535 nm) was recorded using FP-8300 spectrofluorometer (JASCO), every 40 sec. Total measurement time was set to 3240 sec. The excitation bandwidth was fixed to 5 nm, and the emission bandwidth was fixed to 10 nm. As a background control, the reaction sample containing only 100 μM donor liposomes was recorded (“donor only”). The fluorescence increase (ΔEm535 nm) at each time point was calculated by subtracting the NBD signal recorded from the “donor only” group.

### Statistical analysis

Statistical analyses were performed using GraphPad Prism version 9.5.1 for macOS, GraphPad Software, San Diego, California, USA (www.graphpad.com) according to their recommendations. Normality of the data as well as SD similarity were tested before any statistical test for differences was performed. For two sample comparisons, an unpaired t-test was used for Gaussian-distributed data with similar SD, while Welch correction was applied for normal data with different SD distribution. In cases where Gaussian distribution could not be assumed, the Mann-Whitney test for rank comparisons was performed to determine significance. For more than 2 sample comparisons, ordinary One-way ANOVA followed by Dunnett’s multiple comparisons test was used when data followed Gaussian distribution and presented equal variance. In normal-distributed datasets where SD was different, Brown-Forsythe and Welch ANOVA tests were performed followed by Dunnett’s T3 multiple comparisons test. For data that did not pass the normality tests, statistical significance was calculated using the Kruskal-Wallis test followed by Dunn’s multiple comparisons tests. Significance is noted as: ns > 0.05, * p < 0.05, ** p < 0.005, *** p < 0.001, **** p < 0.0001.

## Acknowledgments

The authors thank the following: Jay Xiaojun Tan (Aging Institute, University of Pittsburgh School of Medicine and University of Pittsburgh Medical Center, Pittsburgh, PA, USA) for advice, discussion and sharing reagents and protocols; Andrew Peden (University of Sheffield, UK) for advice and sharing reagents and protocols; Michael Marks (Perelman School of Medicine, University of Pennsylvania, Philadelphia, USA) and Tamas Balla (National Institute of Child Health and Human Development, NIH, Bethesda, USA) for advice and sharing reagents and protocols; Kenji Maeda (Cell Death and Metabolism, Danish Cancer Institute, Copenhagen, Denmark) for helpful advice; Christophe Queval and Michael Howell of the High Throughput Screening Core facility (Crick Institute) for advice and discussions; Rocco D’Antuono of the Light Microscopy Core facility (Crick Institute) for advice and discussions; Mark Skehel of the Proteomics Core facility (Crick Institute) for helpful advice; all members of the Tooze lab for lively discussions.

## Author contributions

Conceptualization: SDT, EA, SAT

Methodology: SDT, EA, DDY, WZ, EM, JHH, EP, IP, DF, TLMT, MG, SAT Investigation: SDT, EA, DDY, WZ, EM, JHH, EP, IP, DF

Visualization: SDT, EA, DDY, WZ, EM, JHH, EP, IP, DF

Funding acquisition: TLMT, MG, SAT Project administration: SDT, EA, SAT Supervision: SDT, EA, TLMT, MG, SAT Writing – original draft: SDT, EA, SAT

Writing – review & editing: SDT, EA, DDY, WZ, EM, JHH, EP, IP, DF, TLMT, MG, SAT

## Data and materials availability

All data are available in the main text or the supplementary materials.

