## Supplementary Figures for "ATG9A and ARFIP2 cooperate to regulate PI4P levels for lysosomal repair"

### SUPPLEMENTARY INFORMATION

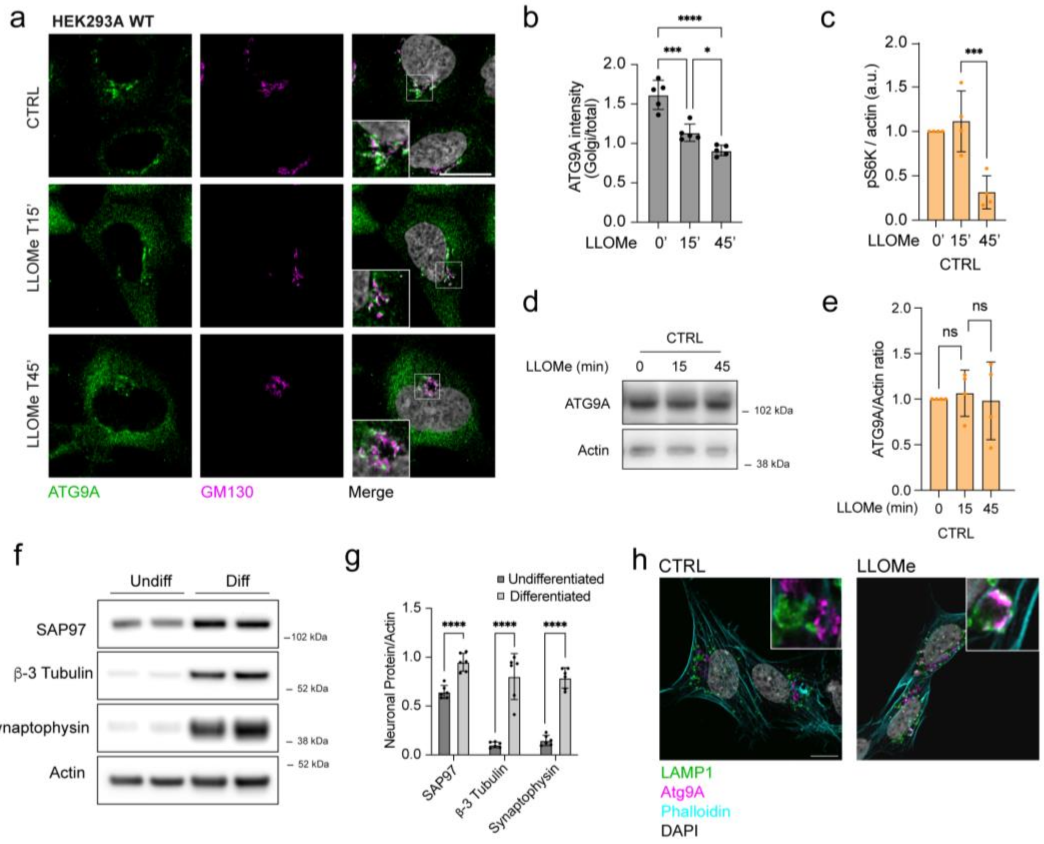


**Extended Data Fig. 1. ATG9A relocalizes on lysosomes upon lysosomal damage.** (a) HEK293A WT cells were treated with 1 mM LLOMe for the indicated times followed by immunofluorescence. Scale bar: 10 µm. (b) Quantification of ATG9A on the Golgi. *n* = 5 independent experiments, * p < 0.05, *** p < 0.001, **** p < 0.0001. (c) Quantification of phospho-S6-Kinase intensity. *n* =65 independent experiments, *** p < 0.001. (d) CTRL cells were treated with 1 mM LLOMe for the indicated times and Western Blot was performed. (e) Quantification of ATG9A intensity from (d). *n* = 4 independent experiments. (f) SH-SY5Y cells were differentiated as described. Undifferentiated (undiff) and differentiated (diff) cells were analyzed by Western Blot and neuronal markers were used. (g) Quantification of (f) shows an increase in differentiation markers. *n* = *6* independent experiments, **** p < 0.0001. (h) Differentiated SH-SY5Y cells were treated with LLOMe 1 mM for 15 minutes and subjected to immunofluorescence analysis. Scale bar: 10 µm.

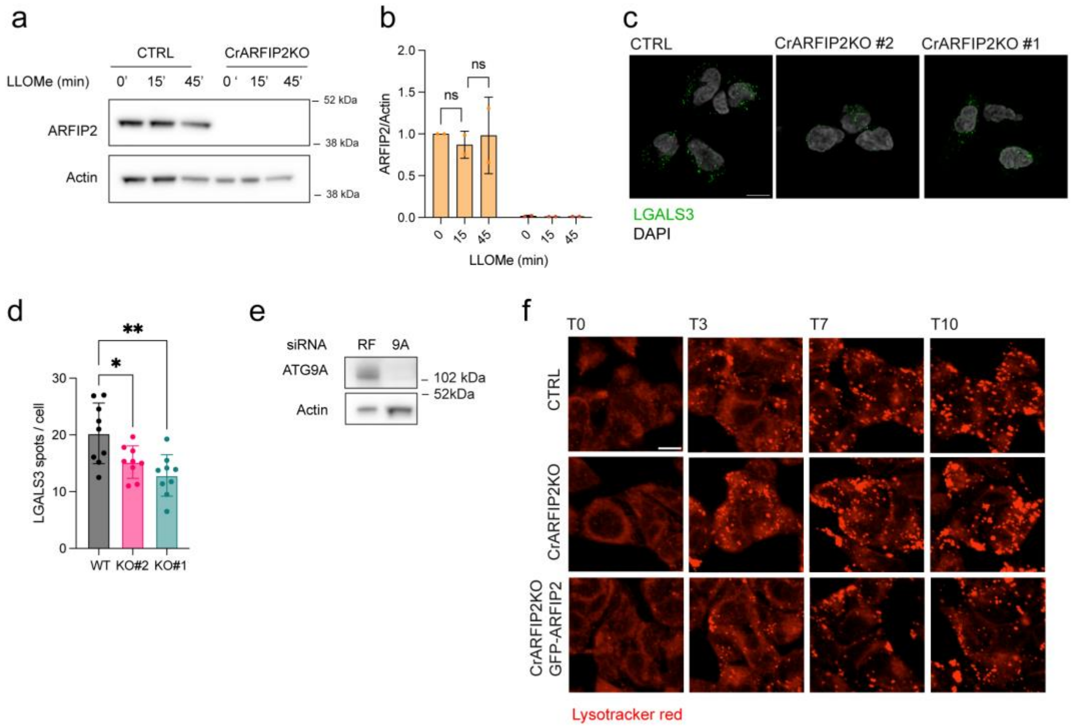


**Extended Data Fig. 2. ARFIP2 regulates lysosomal repair through ATG9A.** (a) HEK293A CTRL and CrARFIP2KO cells were treated with 1 mM LLOMe for the indicated times and ARFIP2 protein levels were analyzed by Western Blot. (b) Quantification of ARFIP2 levels. *n = 2* independent experiments. (e) Western Blot showing depletion of ATG9A upon siRNA. (c) HEK293A CTRL and CrARFIP2 clone 1 and clone 2 were treated with LLOMe 1 mM for 15 minutes followed by immunofluorescence. Scale bar: 10 µm. (d) Quantification of LGALS3 spots per cell was performed. *n=2* independent experiments, * $p < 0.05$ ** $p < 0.01$. (f) Representative images of Lysotracker recovery after washout in HEK293A CTRL, CrARFIP2KO and rescue cell lines at different time point (minutes).

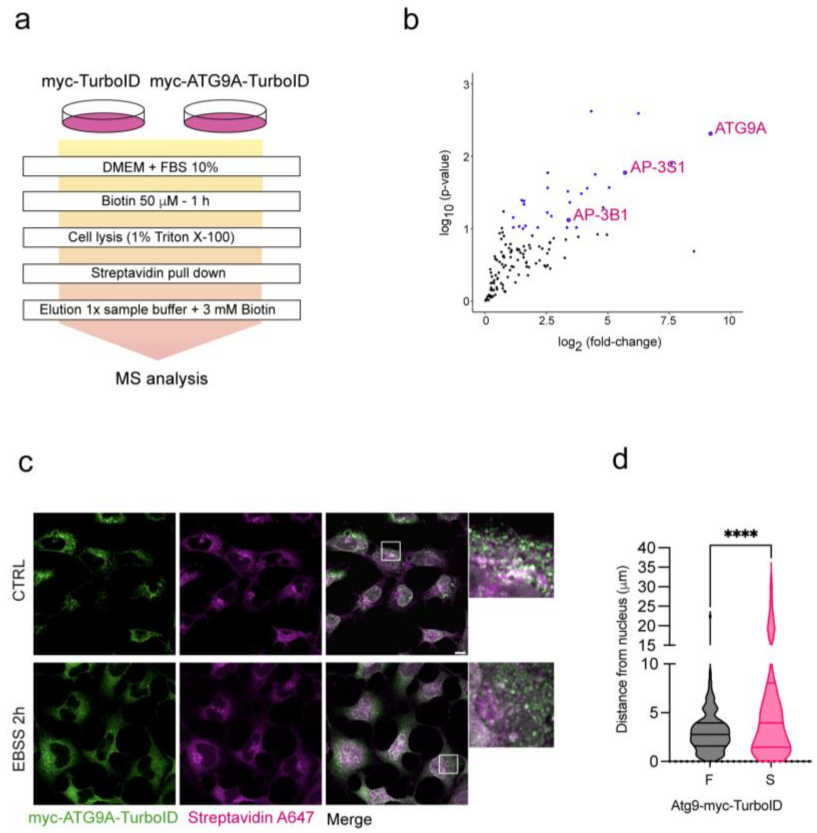


**Extended Data Fig. 3. MS experiments using myc-ATG9A-turbo ID revealed AP-3 as interactor of ATG9A.** (a) Schematics of the mass spectrometry protocol followed to analyze ATG9A proximity proteome. (b) Plot representing ATG9A proximal interactors. Subunits of the AP-3 complex and ATG9A are highlighted. (c) HEK293A expressing myc-ATG9A-TurboID were subjected to starvation (EBSS 2h) and ATG9A dispersal was analyzed by immunofluorescence. Scale bar: 10 µm (d) Quantification of ATG9A dispersal upon starvation. *n* = 3 independent experiments, **** $p < 0.0001$.

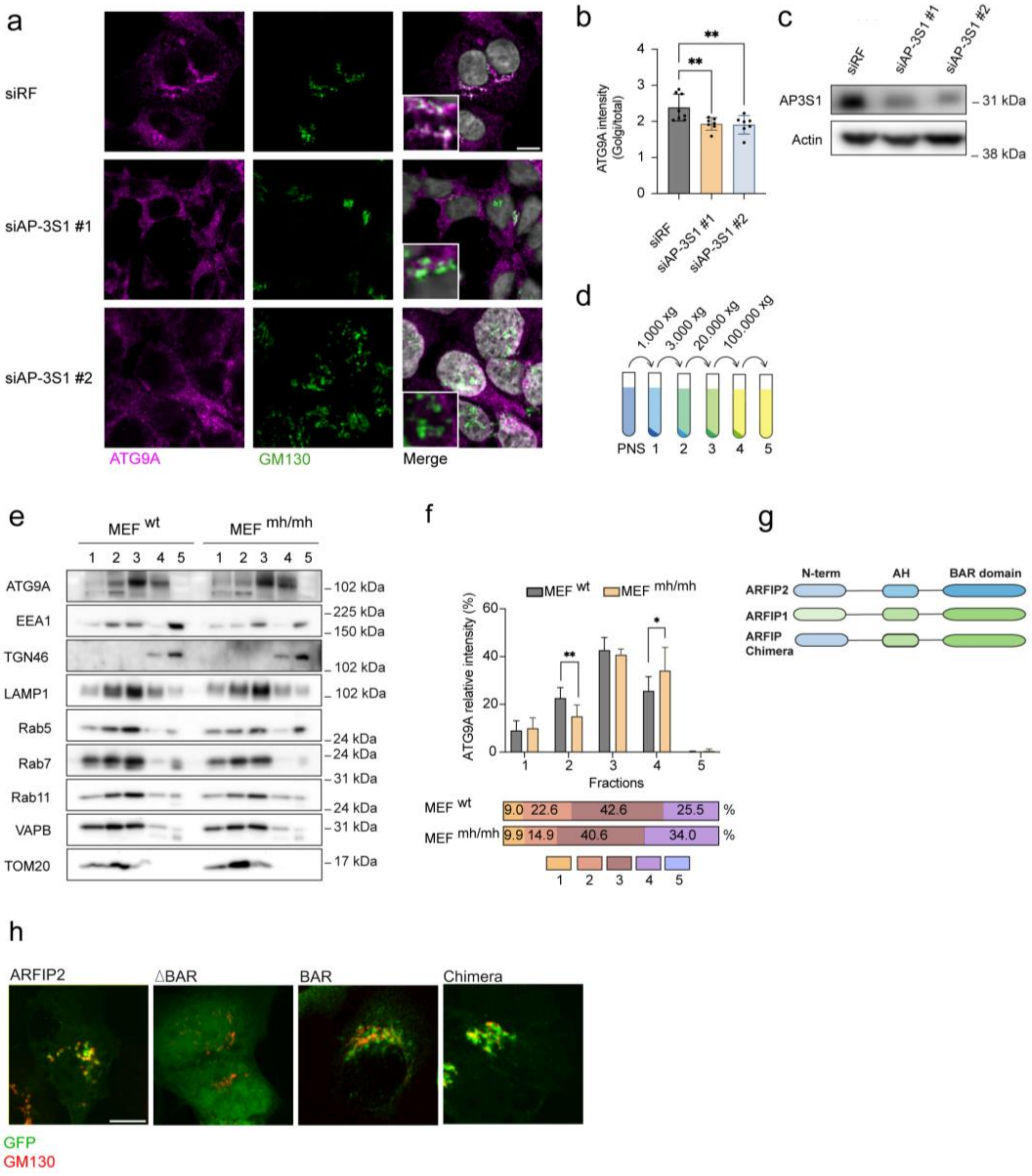


**Extended Data Fig. 4. AP-3 regulates ATG9A trafficking and interacts with ARFIP2 N-terminal domain.** (a) HEK293A WT cells were transfected with siRF or siRNA AP-3S1 subunit (#1 and #2) to deplete AP-3 complex. Immunofluorescence was performed to follow ATG9A localization. Scale bar: 10 μm. (b) Quantification of ATG9A on the Golgi. $n$ = 3 independent experiments, ** $p < 0.01$. (c) Western Blot from HEK293A cells transfected with siRNAs for AP-3S1 subunit. (d) Schematic representing the differential centrifugation protocol used for membrane fractionation. (e) Western Blot analysis of organelle markers and ATG9A in $MEF^{wt}$ and $MEF^{mh/mh}$ from collected pellets. (f) Quantification of the ATG9A proportion in the different fractions. $n$ = 3 independent experiments, * $p < 0.05$, ** $p < 0.01$. (g) Schematic representation of the structural domains of ARFIP2 WT, ARFIP1 WT and ARFIP chimera used in this study. (h) CrARFIP2KO cells transiently expressing GFP-ARFIP2, GFP-deltaAH-BAR (1-108 aa), GFP-AH-BAR (109-341 aa) and GFP-ARFIP Chimera were processed for immunofluorescence. Scale bar: 10 μm.

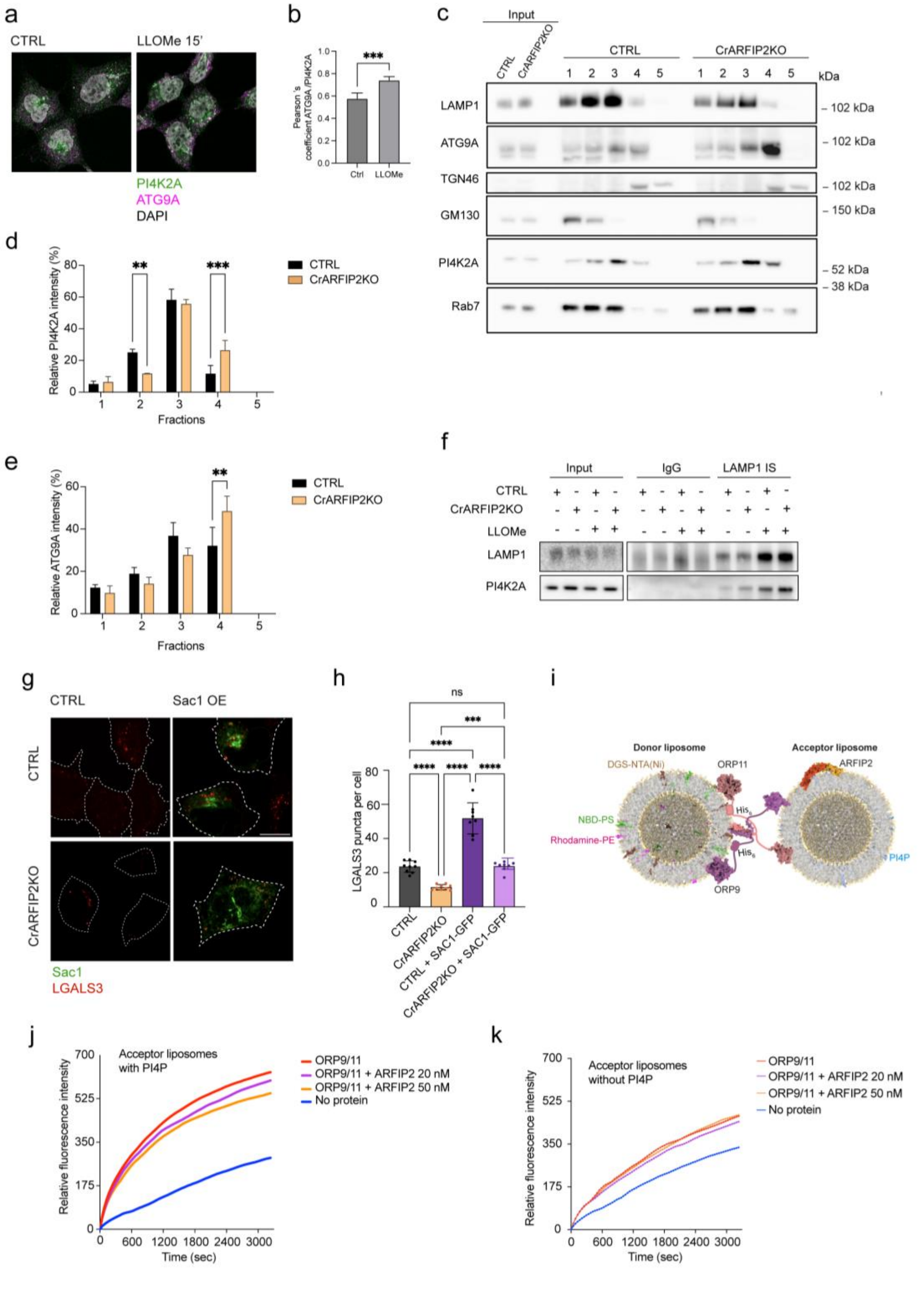


**Extended Data Fig. 5. ARFIP2 mediates lysosomal repair through PI4P and modulates ORPs-dependent lipid transfer.** (a) HEK293A cells were treated with LLOMe for 15 minutes followed by immunofluorescence. Scale bar: 10 µm. (b) Quantification of (a). *n* = 5 fields (100 cells), *** $p < 0.001$. (c) Western Blot from CTRL and CrARFIP2KO cells after membrane fractionation. (d-e) Quantification of the PI4K2A (d) and ATG9A (e) proportion in the different fractions. *n* = 3 independent experiments, ** $p < 0.01$, *** $p < 0.001$. (f) Lysosome purification was performed using an antibody endogenous LAMP1 in CTRL and CrARFIP2KO cells treated with LLOMe for 15 minutes. Western Blot analysis was performed. (g) CTRL and CrARFIP2KO cells were transfected with GFP-Sac1 and subsequently treated with 1 mM LLOMe for 45 minutes. Scale bar: 10 μm (h) Quantification of LGALS3 spots in CTRL and CrARFIP2KO cells after overexpression of Sac1. $n = 3$ independent experiments, *** $p < 0.001$, **** $p < 0.0001$. (i) Schematic model of the *in vitro* lipid transport assay for ORP9/11 heterodimer and ARFIP2. Donor liposomes were enriched with NBD-PS (green), Rhodamine-PE (magenta) and DGS-NTA(Ni) lipids, while acceptor liposomes were enriched or not with PI4P (blue). Both ORP9 and ORP11 contain a $His_6$-tag to bind DGS-NTA(Ni) in the donor liposomes. The structure models for ORP9/11 heterodimer and ARFIP2 homodimer were obtained using AlphaFold/AlphaFold Multimer modelling. (j-k) PS transfer activity of ORP9/11 heterodimer in the presence or absence of ARFIP2 at the reported concentrations using acceptor liposomes with PI4P (j) or without PI4P (k).

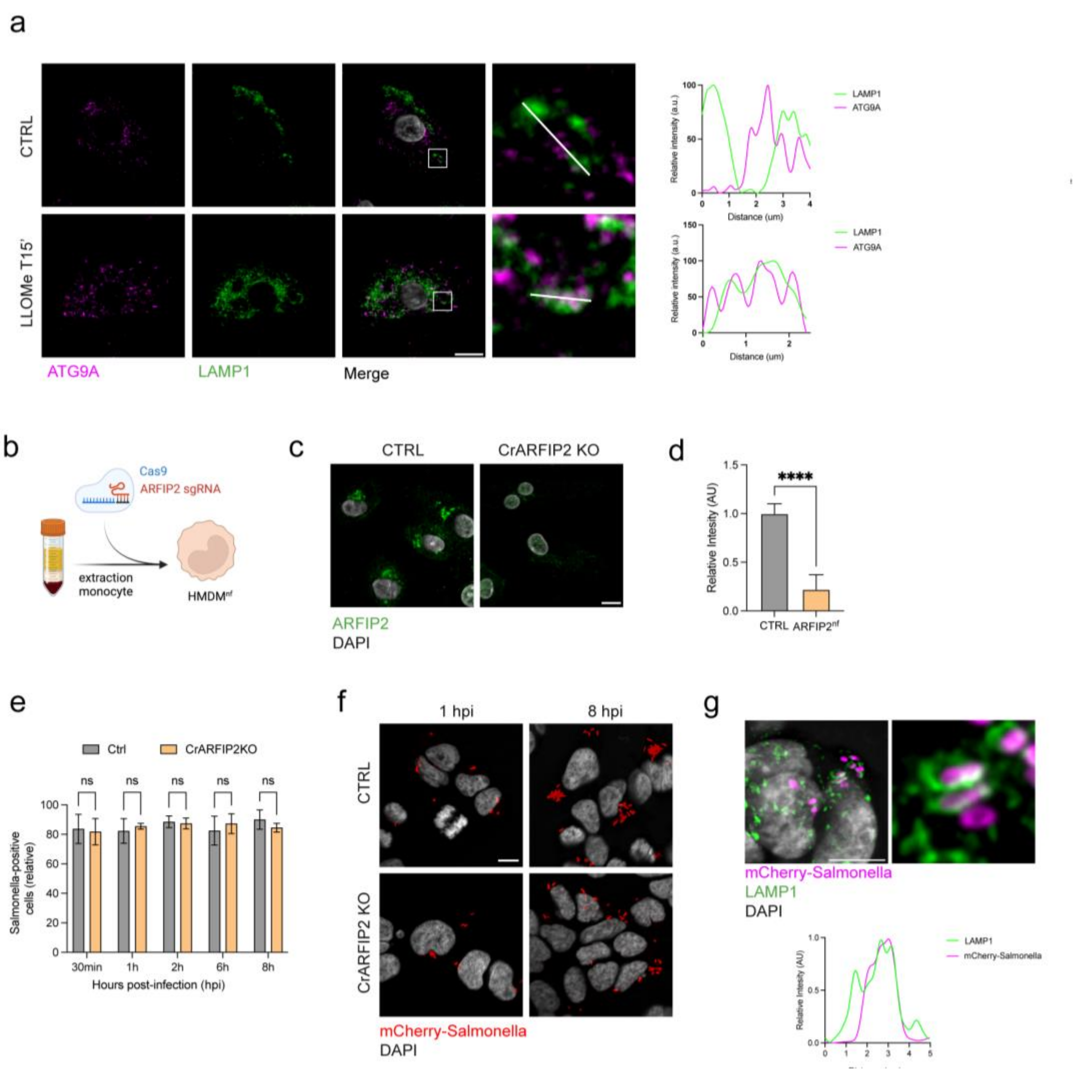


**Extended Data Fig. 6. ARFIP2 regulates bacterial infection through lysosomal repair.** (a) Immunofluorescence of HMDM cells treated or not with 1 mM LLOME for 15 minutes and stained for ATG9A and LAMP1. Scale bar: 10 µm (b) Schematic of the protocol used to generate HMDM cells nucleofected with CRISPR guides. (c) Representative images of CTRL and ARFIP2$^{nf}$ HMDM cells. Scale bar: 10 µm. (d) Quantification of ARFIP2 fluorescence in CTRL and ARFIP2$^{nf}$ HMDM cells. *n* = 3 independent experiments, **** p < 0.0001. (e) Quantification of cells infected with *Salmonella* in CTRL and CrARFIP2KO cells at the indicated time points post-invasion. *n* = 3 independent experiments. (f) Representative images of mCherry-*Salmonella*-infected cells in CTRL and CrARFIP2KO at the indicated hours post-invasion (hpi). Scale bar: 10 µm. (g) Immunofluorescence analysis of *Salmonella* and LAMP1. (top) A line plot was used to represent bacteria engulfed in LAMP1-positive compartment after 30 min post-invasion (bottom).
